# Hoisted by Their Own Petard: Defeating Immunity Protein–Mediated Bacteriocin Resistance

**DOI:** 10.1101/2025.10.23.684137

**Authors:** Tim Moritz Weber, Raquel Guadarrama-Gonzalez, Colin R. MacKenzie, Jörg Pietruszka

## Abstract

The irresponsible use of antibiotics has led to an erratic spread of antimicrobial resistance (AMR) among bacterial pathogens. Without intervention, 10 million AMR-related deaths per year by 2050 are estimated by the WHO. Nonetheless, the development of novel antibiotics has almost come to a halt. In this context, naturally occurring narrow-spectrum antimicrobial proteins, known as bacteriocins, are promising alternatives to address the need for effective and selective antimicrobial treatments. However, the widespread distribution of cognate immunity proteins among bacteriocin producers vastly limits the natural antimicrobial spectrum against Gram-negative bacteria. On the example of pyocin S2 (PyoS2), an S-type HNH DNase bacteriocin targeting the opportunistic human pathogen *Pseudomonas aeruginosa*, this study demonstrates for the first time that a rationally introduced single mutation and chemical protein functionalisation at PyoS2’s immunity protein exosite fully eradicate immunity protein-mediated resistance to pyocins in multidrug-resistant clinical *P. aeruginosa* isolates. The presented findings highlight a strategy to engineer bacteriocins from pathogenic Gram-negative bacteria with an extended antimicrobial spectrum, overruling the prevalent and predominant bacteriocin resistance mechanism of immunity protein binding. This strategy could expedite the development of engineered bacteriocins to expand our repository of antimicrobials to combat the worsening antimicrobial resistance crisis and avert a post-antibiotic era.

## Introduction

Antimicrobial resistance (AMR) to last-resort antimicrobials is one of our society’s most urgent medical challenges. If no action is taken now, the number of deaths attributable to and associated with AMR will increase to 10 million deaths by 2050^[1]^. While the discovery of new antibiotics and cellular bacterial targets is stagnating, the rapid spread of AMR among pathogenic bacterial species in hospitals is of most significant concern. Therefore, the demand for effective antimicrobial agents is globally high, calling for innovative approaches to mine for novel antimicrobials and fulfil the need for these indispensable therapeutics^[2]^.

A genuine alternative to antibiotics are bacteriocins^[3]^, natural antimicrobial proteins produced by bacteria of all kinds for intraspecies competition^[4-5]^. The bacteriocins of Gram-positive species often possess a broad antimicrobial spectrum and find wide application in food industries for preservation (e.g. nisin or enterocins)^[6-7]^. In contrast, the bacteriocins of Gram-negative organisms surprisingly still lack proof of applicability so far^[8]^. Unlike most antibiotics, Gram-negative bacteriocins are narrow-spectrum antimicrobials, typically killing only closely related species within the same genus (e.g., colicins from *Escherichia coli* target *E. coli*)^[9-11]^. This limited activity spectrum is predominantly shaped by three elements: the highly variable architecture of the outer membrane (outer membrane receptors and cell surface lipopolysaccharides), the strain-selectivity of energy-transducing complexes (Ton and Tol) used for bacteriocin membrane translocation, and a widespread prevalence of cognate immunity proteins among bacteriocin producers. Although the narrow activity spectrum in conjunction with distinct cytotoxic mechanisms from common antibiotics makes bacteriocins auspicious candidates for combating AMR, the widespread distribution of natural immunity proteins conflicts with their intended use as antimicrobial agents. Hence, overcoming the protective mechanism of immunity protein binding is required to expand the natural activity spectrum of bacteriocins.

One of the major contenders in humanity’s race against AMR is the opportunistic human pathogen *Pseudomonas aeruginosa*, which has evolved elaborate escape and virulence mechanisms (e.g., biofilms, efflux pumps, and antibiotic-hydrolysing enzymes). These mechanisms enable the Gram-negative bacterium to establish persistent nosocomial infections in immunocompromised patients, making this bacterium extremely difficult to treat^[12-13]^. Moreover, the spread of transferable carbapenem resistance among clinical strains of *P. aeruginosa* has been recognised as a significant safety risk in hospitals, underscored by the high mortality associated with infections caused by these strains^[14-15]^. Accounting for the growing threat of AMR caused by transferable carbapenemase genes, the World Health Organization (WHO) published in 2017 a priority list of bacterial pathogens for which new antibiotics or alternative therapeutics are urgently needed, ranking carbapenem-resistant *P. aeruginosa* (CRPsA) in the highest priority category “critical”^[2, 16]^.

Although severe nosocomial infections with CRPsA already require tailored antimicrobials for effective treatment and infection control, the pathogen also provides us with possible alternatives to those precious antibiotics. More than 90% of all known *P. aeruginosa* strains are known to produce different classes of bacteriocins (pyocins)^[17]^. Of the 17 experimentally validated or predicted pyocins belonging to the class of soluble S-type pyocins (Supplementary Table 1), most hijack the conserved TonB machinery and the essential inner membrane (IM) AAA+ protease FtsH for membrane translocation^[18-19]^. The currently best-studied S-type pyocin is the class III HNH DNase bacteriocin pyocin S2 (PyoS2). On the cell surface, PyoS2 first binds to the common polysaccharide antigen (CPA), before hijacking the TonB-dependent ferripyoverdine siderophore receptor FpvAI for outer membrane (OM) translocation^[20-21]^. Upon FtsH-catalysed processing at the inner membrane, the cytotoxic C-terminal DNase domain is guided into the target cell’s cytoplasm, where the PyoS2 DNase unleashes cytotoxic activity by degrading the host cell’s DNA^[22]^. This DNA degradation then triggers the SOS response, and ultimately leads to cell death (Supplementary Fig. 1a). However, strains producing PyoS2 protect themselves from their own toxin by expressing the cognate immunity protein S2 (ImS2), which forms a stoichiometric neutralising complex by binding with femtomolar affinity to the immunity protein exosite (IPE) within the toxin’s cytotoxic domain^[23-24]^. This toxin-antitoxin relationship, common to all S-type pyocins and most other bacteriocins from Gram-negative bacteria, vastly limits the antimicrobial spectrum of natural bacteriocins and thereby limits their impact as potential therapeutics.

So far, little effort has been made to translate our current knowledge on Gram-negative bacteriocins into application, harnessing their antimicrobial potential for combating infections with MDR strains. In this work, we challenge the principle of immunity protein-mediated bacteriocin resistance, a prerequisite for developing bacteriocins of Gram-negative bacteria as viable next-generation antimicrobials. On the example of S-type PyoS2 and its cognate immunity protein ImS2, this study provides evidence for the involvement of the O serotype in S-type pyocin susceptibility. Additionally, this case study devises a general strategy to disrupt the ultra-high-affinity interaction between toxin and immunity protein through rational protein engineering and chemical modification, collectively overruling the inherent ImS2-dependent PyoS2 resistance of CRPsA clinical isolates.

## Results

### A few residues mediate ultra-high-affinity binding between PyoS2 and ImS2

Pyocin S2 is a multi-domain protein, for which no experimental full-length structure has been reported to date. However, structure predictions have been made available by AlphaFold (AlphaFold DB AF-Q06584-F1-v4). Together with experimental findings from other bacteriocin homologues, the structural model reveals that the 74-kDa protein comprises two tandemly repeated N-terminal kinked three-helix bundle domains (Fig. 1a). The N-terminal bundle is required for binding to the FpvAI primary OM receptor and translocations through its lumen via the TonB-dependent pathway (residues 50−195, cf. Supplementary Fig. 1a)^[25]^. The second bundle is used for binding to the CPA lipopolysaccharide on the cell surface (residues 196−320)^[26]^, enhancing the overall affinity to the bacterial membrane. The C-terminal segment of PyoS2 is segregated into two domains, one domain required for FtsH processing and IM translocation (residues 320−556)^[27]^, and the C-terminal cytotoxic domain harbouring the HNH nuclease (residues 557−689)^[24]^. The latter forms a stoichiometric neutralising complex with the cognate 10-kDa immunity protein ImS2. Other than full-length PyoS2, the DNase/ImS2 complex has already been crystallised at 1.8 Å resolution (PDB-REDO 4QKO)^[24]^ (Fig. 1a2), showing a high congruency for the DNase domain between the AlphaFold model and the crystal structure (RMSD 0.69 Å). Although HNH DNase bacteriocins and their cognate immunity proteins associate with femtomolar affinity^[28]^, the strong binding can be broken down into a handful of interacting amino acids. A comparison of HNH DNase domains from multiple species also reveals that the sequence portion representing the IPE harbours the least sequence conservation, as this portion of the DNase domain mediates high-affinity binding to the sequentially unique cognate immunity proteins (Fig. 1b).

**Fig. 1.**
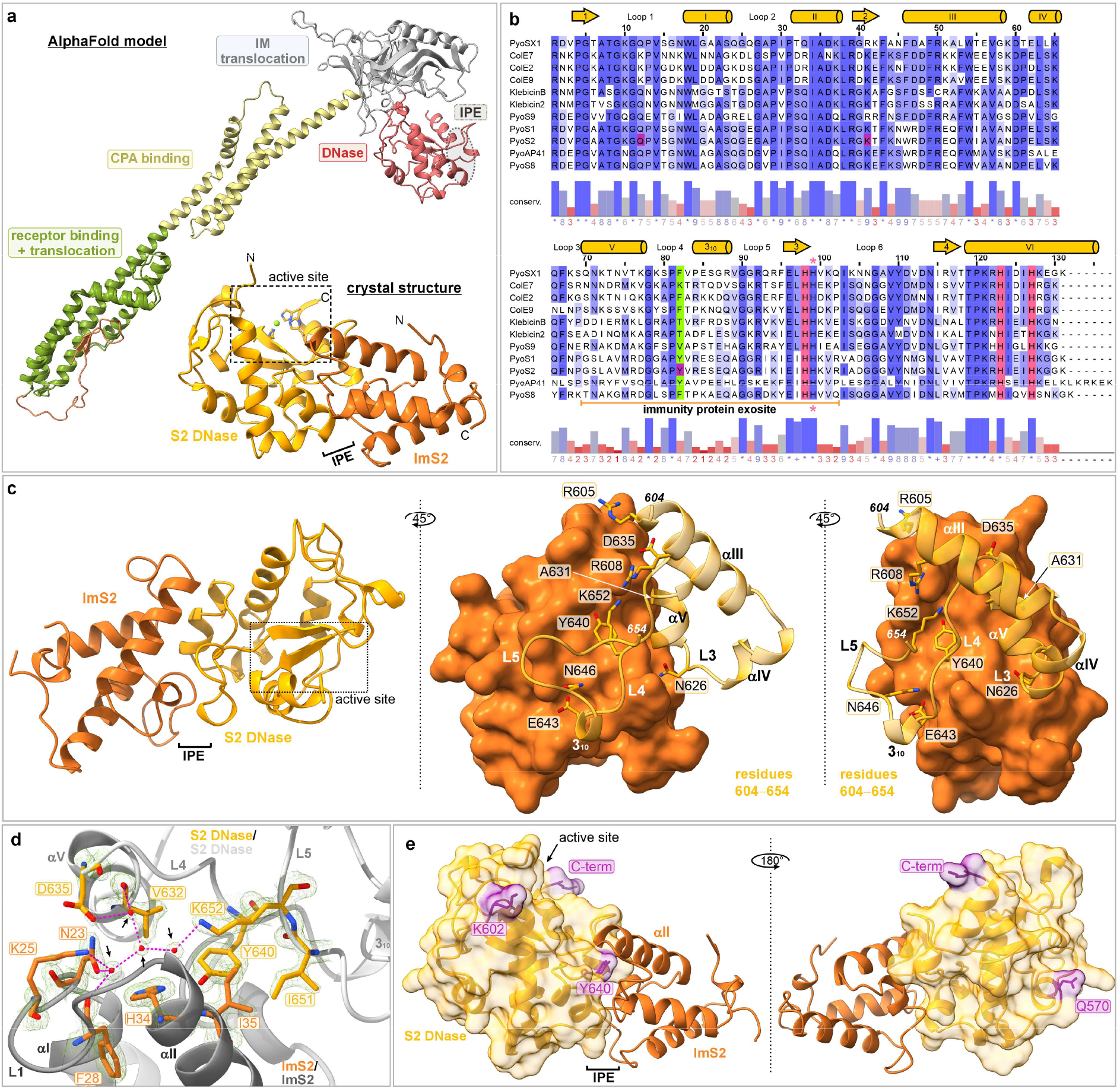
Structural analysis of PyoS2 and its cognate binding to ImS2. **a**. AlphaFold model of PyoS2 (AF-Q06584-F1-v4) without ImS2 and crystal structure of the PyoS2/ImS2 complex (PDB-REDO 4QKO). **b**. Multiple sequence alignment of DNase domains of HNH DNase bacteriocins. Positions equivalent to Y640 in PyoS2 are highlighted in green. Conserved histidine residues required for DNase activity are highlighted in red (metal complexation) or marked with a pink asterisk (general base). Residues selected for cysteine substitutions are highlighted in purple. **c**. Involved PyoS2 IPE residues in ultra-high-affinity binding to ImS2. **d**. Network of conserved interstitial waters at the PyoS2 IPE. The crystal structure is aligned with the 2Fo-Fc electron density map (green mesh) at 0.6σ in a range from −0.836 − 5.16. Shown are the interacting residues (PyoS2 DNase in yellow, ImS2 in orange) and the protein backbones (PyoS2 DNase in white, ImS2 in grey). Waters are depicted as red spheres (highlighted by black arrows). **e**. DNase residues selected for cysteine substitutions and addition, and their relative position to ImS2 and the DNase’s active site.

Accordingly, we hypothesise that knowledge of those residues required for neutralising complex formation and mediating binding selectivity can be used to engineer bacteriocins that evade neutralisation of DNase activity through immunity protein binding.

Based on molecular interface analyses of the PyoS2/ImS2 crystal complex with PDBePISA^[29]^, we identified nine amino acids within the IPE of PyoS2 that show attraction to ImS2 via hydrogen bonding or salt bridges (Supplementary Table 2, Fig. 1c). From the identified residues R605, R608 (positioned in helix III), N626, A631 (helix V), D635, Y640 (loop 4), E643, N646 (3_10_ helix), and K652 (loop 5) (Fig. 1b and 1c), D635 and Y640 are located in the large loop 4, which is devoid of any secondary structure. Additionally, both residues are enclosing a stabilising network of four interstitial water molecules that is found at the interface of PyoS2/ImS2 and is present within all four asymmetric units of the crystal unit cell (Supplementary Fig. 1b). This network with well-defined spherical electron density in the 2Fo-Fc density map (Fig. 1d) is adjuvant in intraprotein stabilisation within both, the DNase and ImS2, and mediating interprotein contacts within the PyoS2/ImS2 complex. For example, the water network enhances the binding of the PyoS2/ImS2 complex by fixing the distorted orientation of ImS2 loop 1, which is required for hydrogen bond formation between the ImS2 residues N23 and K25 with the DNase loop 4 residue D635. The D635 residue also interacts with K652 through the extended water network, stabilising the orientation of the ε-amino group at the interface. Additionally, the interface is stabilised by strong polar interactions and salt bridges between Y640 and K652 with the ImS2 residues H34 and E31 (cf. Supplementary Fig. 1b), respectively, both located in the energy binding hotspot helix II of ImS2.

Because of the involvement of Y640 in stabilising the interstitial water network and its exposed positioning at the IPE, leading to direct interaction with ImS2, we hypothesised that this residue takes a pivotal role in rendering ultra-high-affinity binding and protecting PyoS2-producing strains from self-inflicted death. Together with the relatively small binding interface between the two proteins, we sought to introduce mutational changes and install bulkier non-proteinogenic labels at the IPE of PyoS2 to overcome ImS2-dependent PyoS2 resistance of *P. aeruginosa*.

### Structural disruption at PyoS2’s IPE undermines immunity protein-mediated protection of DNA

Because of the close positioning of the DNase loop 4 residue Y640 next to the stabilising network of interstitial water molecules at the PyoS2/ImS2 interface and its exposed orientation mediating direct polar interactions with ImS2, we decided to substitute residue Y640 with a cysteine residue. This small nucleophilic amino acid facilitates further site-selective functionalisation via maleimide chemistry (Supplementary Fig. 2f), thereby putatively blocking the PyoS2/ImS2 complex association. The chemical functionalization of cysteines with maleimides (mostly *N*-ethylmaleimide) is a rarely used strategy to mitigate, or even prevent, the association of protein complexes through generating a physical barrier at the protein-protein interface^[30-32]^. Besides position Y640 within PyoS2’s IPE, three further cysteine residues were also installed in unstructured regions distant from the IPE. Those residues were placed in the DNase loop positions Q570C (loop 1) and K602C (unstructured region between β-strand 3 and helix III), as well as through a cysteine addition in the C-terminal position (C-Cys) at the extension of helix VI (cf. Fig. 1e). Alongside cysteine mutants, DNase inactive PyoS2 variants were generated through the substitution of the mechanistically essential “general base” residue H657A and the metal-coordinating residue H685 with alanine residues. Both histidine residues are highly conserved among HNH DNase bacteriocins (cf. Fig. 1b) and are obligatory for efficient HNH DNase activity.

All proteins were heterologously expressed in a stoichiometric complex with ImS2 in *E. coli* BL21(DE3) (Supplementary Fig. 2a). However, for the PyoS2 Y640C mutant, no reliable expression could be established (Supplementary Fig. 2a and 2b), hinting toward a toxic phenotype. This toxicity was assumed to be caused by insufficient complexation of PyoS2 by ImS2, resulting in the degradation of DNA within the host cell. Subsequently, *E. coli* strains JM109(DE3)^[33]^, Rosetta 2(DE3) pLysS, and C43(DE3)^[34]^, common hosts for the expression of toxic proteins, were screened for their potential as suitable expression hosts for PyoS2 Y640C (Supplementary Fig. 2c), leading to the identification of strain C43(DE3) as an ideal host for the production of this PyoS2 mutant. In the subsequent downstream processing, PyoS2 variants were purified using a two-step chromatographic protocol, yielding highly pure proteins (Supplementary Fig. 2d; validated by mass spectrometry: Supplementary Table 3).

To fathom the ability of ImS2 to neutralise the DNase activity of PyoS2 and its generated mutants and conjugates to rescue plasmid DNA from degradation, the purified PyoS2 variants were next subjected to a plasmid nicking assay^[35]^. In this regard, a single time point analysis was performed after 15 h for all PyoS2 mutants and conjugates (Fig. 2a and 2b), complemented by kinetic investigations for selected DNase active variants for the determination of supercoiled DNA half-life (DNA_S_ *t*_1/2_) in the presence and absence of ImS2 (Fig. 2c, Supplementary Fig. 3, 4, and 5, Supplementary Table 4, 5, and 6).

**Fig. 2.**
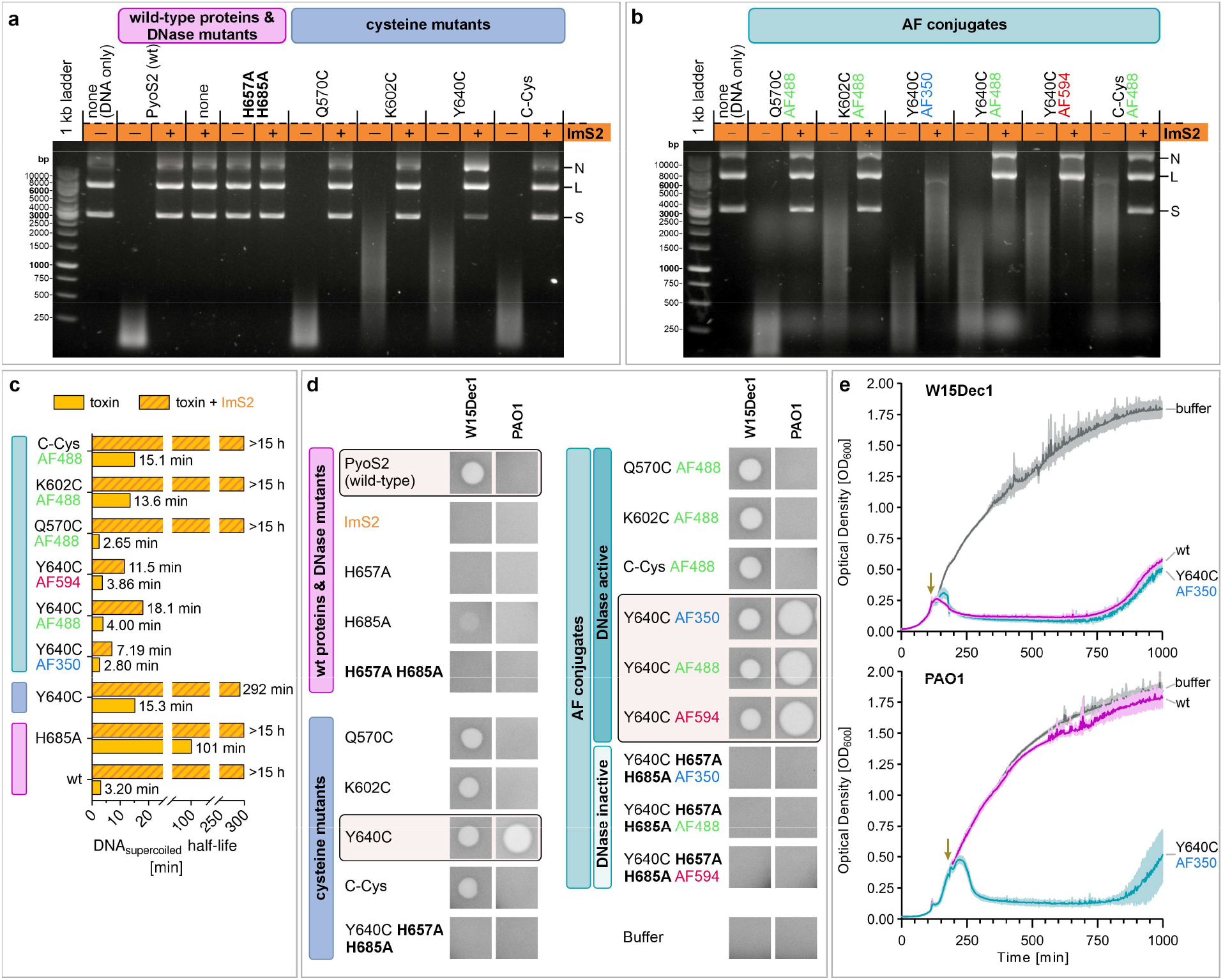
Structure alteration at the IPE prevents effective ImS2 binding and overrules PyoS2 resistance of strain PAO1. **a**. Plasmid nicking assay with PyoS2 mutants in the presence (+) or absence (−) of ImS2. The plasmid forms nicked (N), linear (L), and supercoiled (S) are annotated accordingly. Recurring colour code: wt proteins and DNase mutants (pink), cysteine mutants (blue), AF conjugates (green). **b**. Plasmid nicking assay with PyoS2-AF conjugates in the presence or absence of ImS2. **c**. Half-life of supercoiled DNA for selected DNase active PyoS2 mutants and conjugates in the presence and absence of ImS2. **d**. Plate killing assay with strains W15Dec1 (PyoS2-sensitive) and PAO1 (PyoS2-resistant) and several PyoS2 mutants and conjugates at 5 µM. Clear spots indicate antimicrobial activity. **e**. Turbidity assay in LB media with PyoS2 (wt) and the PyoS2 Y640C-AF350 conjugate on strain PAO1 (PyoS2-resistant) and W15Dec1 (PyoS2-sensitive). Small arrows indicate toxin addition at 750 nM after 1:49 h (W15Dec1) or 2:51 h (PAO1).

Starting with wild-type PyoS2 as a positive control, the DNase active toxin had fully digested all three species of the DNA substrate (nicked, linear, supercoiled) after 15 h through unspecific hydrolysis. This resulted in low-molecular weight DNA fragments (<350 bp) (Fig. 2a) and was characterised by a DNA_S_ half-life of 3.20 min (Fig. 2c). In contrast, the addition of ImS2 in slight excess (5 µM PyoS2 vs. 5.25 µM ImS2) thoroughly protected the DNA from PyoS2-catalysed degradation, leaving all three DNA species unharmed (*t*_1/2_ >15 h). No catalytic DNase activity was observed instead for ImS2 alone. Also, the PyoS2 H657A H685A double mutant showed no residual activity due to the removal of the general base residue H657 and the metal-coordinating residue H685 located in helix VI (Supplementary Fig. 6a), both of which are required for efficient HNH DNase activity (negative control; referred to as DNase knock-out). All generated PyoS2 single cysteine mutants (Q570C, K602C, Y640C, and C-Cys) retained DNase activity. However, only Q570C and C-Cys retained activity at a similar level to wild-type PyoS2 (<500 bp with and <900 bp after 15 h, respectively). The mutants K602C and Y640C instead revealed a broader size distribution of remaining DNA fragments after 15 h (150−3000 bp, *t*_1/2_ 15.3 min for Y640C), indicating adverse effects of the single amino acid substitution on DNA binding or catalytic DNase activity. When the immunity protein was added, the DNase activity of PyoS2 variants Q570C, K602C, and C-Cys was neutralised entirely. However, the Y640C mutant retained partial activity even in the presence of ImS2, as visualised by the diminished supercoiled DNA species and the pronounced accumulation of a nicked DNA form after 15 h (*t*_1/2_ 292 min). Collectively, only the introduced Y640C mutation located within the IPE was able to retain PyoS2’s DNase activity in the presence of ImS2, even though the presence of the immunity protein resulted in a drastically increased half-life of DNA_S_ in comparison to Y640C alone (19-fold increase).

In addition to mutational substitutions replacing the surface-accessible native amino acids Q570, K602, and Y640 with the small nucleophilic cysteine residue, the bulky and maleimide-functionalised AlphaFluor488 (AF488) fluorophore dye was conjugated to the DNase cysteine mutants via click chemistry in a thiol-Michael addition (Supplementary Fig. 2g). The fluorophore could potentially provide a physical barrier mitigating or preventing the association of the PyoS2/ImS2 ultra-high-affinity complex and thereby cause insufficient protection of dsDNA. From all four conjugates, Q570C-AF488 retained the highest DNase activity in the absence of ImS2 (<500 bp, *t*_1/2_ 2.65 min) and showed no disadvantageous effect of fluorophore labelling on DNase activity in comparison to the unlabelled Q570C mutant (Fig. 2b). Of all selected residues, Q570 is the only one neither being closely located to the active site nor the bound DNA substrate (cf. Fig. 1e). Also, the K602C-AF488 conjugate exhibited comparable DNase activity to the unlabelled K602C mutant (250-6000 bp, *t*_1/2_ 13.6 min). The C-Cys-AF488 conjugate instead demonstrated a notable decrease in DNase activity (750−10000 bp, *t*_1/2_ 15.1 min), resulting from fluorophore attachment. This observation is attributed to the interference of the fluorophore with binding and release of the DNA substrate at the active site, as the introduced cysteine modification is protruding from helix VI of the DNase. This helix additionally harbours the two active site histidine residues H681 and H685 required for chelation of the essential divalent Mg^2+^ metal cofactor (Supplementary Fig. 6a). However, when ImS2 was added to the three conjugates Q570C-AF488, K602C-AF488, and C-Cys-AF488, again, no residual DNase activity was detected (*t*_1/2_ >15 h), as earlier observed for the corresponding cysteine mutants without fluorophore attachment. When shifting the focus to the Y640C-AF488 conjugate that carries the fluorophore label at the IPE, no apparent change in DNase activity between the labelled conjugate and the unlabelled mutant was detected in the absence of ImS2 (150−3000 bp, *t*_1/2_ 4.00 min). However, when ImS2 was present, the supercoiled DNA was still fully digested by Y640C-AF488 within the observation period of 15 h, thereby highlighting an increased mismatch of the PyoS2/ImS2 interface in comparison to the fluorophore-free mutant Y640C. This mismatch originates from the attachment of the bulky fluorophore, which facilitates a higher degree of retained DNase activity in the presence of ImS2 (*t*_1/2_ 18.1 min vs. 292 min).

To further analyse the effect of label sizes on ImS2 binding at the IPE of PyoS2, the PyoS2 Y640C-AF350 and Y640C-AF594 conjugates with variable size in conjugated π-systems were prepared in addition to the earlier introduced AF488 dye conjugate (Supplementary Fig. 2g). Generally, the Y640-AF conjugates followed the trend that smaller label sizes come with higher DNase activity (AF350 > AF488 ≈ unlabelled Y640C > AF594). Although the Y640C-AF350 conjugate with the attached bulky fluorophore being more active than the unlabelled Y640C mutant seems counterintuitive at first glance, the maleimide labelling partially resembles the original Y640 residue and seems to restore important amino acid side chain contacts within the PyoS2 DNase domain that influence DNase activity and were lost upon introduction of the Y640C substitution (Supplementary Fig. 6c). Within the IPE, the hydrophobic side chains of V632 and I652 flank Y640 and engage in lateral hydrophobic interactions with this aromatic residue, while K652 further stabilises Y640 through axial hydrophobic contacts. Although Y640 is positioned within PyoS2’s DNase domain and is therefore distal from the active site, the interaction with K652 appears mutually stabilising and obligatory for effective DNase activity. Residue K652 not only supports resting of the dsDNA substrate on the meandering DNase loop 5 but also forms a salt bridge between its polar ε-amino group and the negatively charged DNA backbone (Supplementary Fig. 1c). Although Y640 does not directly interact with the DNA substrate, the residue is located in close proximity to the DNA backbone (Supplementary Fig. 6b), causing a clash of bulky labels with DNA binding and resulting in the observed decrease DNase activity with increased label sizes.

### Defective PyoS2/ImS2 association overrules ImS2-mediated PyoS2 resistance of PAO1

After finding that the Y640C mutant and its conjugates retain *in vitro* DNase activity in the presence of ImS2, the antimicrobial activity of the PyoS2 mutants and conjugates against established *P. aeruginosa* model strains was assessed. For this analysis, the strain PAO1 and the environmental *P. aeruginosa* isolate W15Dec1 were selected and subjected to PyoS2 treatment in a soft-agar overlay spot assay^[36-37]^. Both strains genomically encode the FpvAI OM siderophore receptor, which is required for PyoS2 uptake, and are therefore potentially sensitive to treatment with PyoS2 (cf. Supplementary Fig. 1a). However, strain PAO1 is a PyoS2/ImS2 producer itself and is thus resistant to treatment with the wild-type toxin. At the same time, the genome of W15Dec1 lacks the corresponding *pyoS2*_*imS2* locus and is literature-known to possess a PyoS2-sensitive phenotype^[20]^. A toxin concentration of 5 µM was selected in the first place to reveal cytotoxic effects of less active PyoS2 variants as well.

Meeting expectations, wild-type PyoS2 showed killing activity against strain W15Dec1, but left strain PAO1 unharmed for the reasons mentioned above (Fig. 2d). Also, the immunity protein ImS2 and the PyoS2 DNase knock-out mutant H657A H685A, both lacking enzymatic DNase activity, did not inhibit bacterial growth for any of the tested strains. When investigating the unlabelled PyoS2 mutants Q570C, K602C, Y640C, and C-Cys, carrying cysteine residues in flexible loop positions within the DNase domain, all point mutants retained potent killing activity against W15Dec1, proving that the cysteine mutations neither interfere with toxin uptake nor membrane translocation. The AF488 fluorophore attachment to residues Q570C, K602C, Y640C, and C-Cys of the PyoS2 DNase also triggered cell death of strain W15Dec1, providing proof of successful translocation of all PyoS2-AF488 conjugates across the bacterial two-membrane cell wall. In contrast, the AF488 DNase knock-out conjugates, additionally carrying the H657A H685A active site mutations (cf. Supplementary Fig. 6a), lost their antimicrobial activity. The loss of antimicrobial activity of conjugates upon knocking out the DNase activity evidences that the conjugated fluorophores display no cytotoxic activity towards *P. aeruginosa* on their own and that observed killing activity is solely owing to PyoS2’s DNase activity.

In contrast to the PyoS2-sensitive strain W15Dec1, the PyoS2-resistant strain PAO1 effectively neutralised the DNase activity of the Q570C, K602C, and C-Cys mutants and their AF488 conjugates by intracellular expression of the ImS2 immunity protein and their enmeshment in the ultra-high-affinity complex (Fig. 2d). Only the Y640C mutant, which harbours the point mutation at the PyoS2/ImS2 binding interface, rendered killing activity against the otherwise PyoS2-resistant PyoS2 producer. This newly acquired antimicrobial activity of PyoS2 Y640C against the ImS2 producer PAO1 was characterised by larger zones of growth inhibition than previously observed for W15Dec1, and it was further enhanced by the attachment of AF fluorophores to the IPE residue Y640C.

An equivalent effect of overruled ImS2-mediated PyoS2 resistance of strain PAO1 was also observed in liquid culture with planktonic bacteria (Fig. 2e). Single-dose treatment of strain W15Dec1 with wild-type PyoS2 at an OD_600_ of 0.20 caused further bacterial growth to OD_600_ 0.25 (23 min delay), until the bacterial density dropped to 0.11. With the PyoS2 Y640C-AF350 conjugate, treatment at OD_600_ 0.20 was accompanied by a longer delay (49 min) until antibacterial activity was observable, allowing the bacteria to grow to an OD_600_ of 0.30. Here, the decreased DNase activity of the conjugate compared with wild-type PyoS2 most likely accounts for the observed delay in antibacterial activity (cf. Fig. 2a and 2b).

The addition of wild-type PyoS2 to the PyoS2-resistant strain PAO1 at an OD_600_ of 0.35 had the same effect as the addition of buffer, leading to unimpaired growth behaviour and a final OD_600_ of ∼1.8 under both conditions. Only the novel Y640C-AF350 conjugate, with a negatively affected association between the toxin and ImS2, was able to harm strain PAO1, which grew to an OD_600_ of 0.47 (46 min delay) before declining in bacterial density. Interestingly, both tested strains (W15Dec1 and PAO1), which had been previously treated with a single dose of pyocin, were able to recover fitness after 800 min, emphasising the requirement for repetitive and prolonged antibacterial treatment in highly dynamic systems to eradicate pathogenic bacteria. In summary, the reported gain of antimicrobial activity for the PyoS2 Y640C mutant and its conjugates against a PyoS2-resistant strain is the first proof of extended-spectrum bacteriocins that overcome immunity protein-mediated resistance through structural changes at the IPE.

### IPE mutations abolish PyoS2 resistance of MDR isolates and reveal a serotype-dependent component of S-type pyocin killing

To appraise the significance of our finding that small structural changes at the IPE overrule the ImS2-mediated PyoS2 resistance, we set out to test the generated PyoS2 derivatives on clinical CRPsA strains, which were isolated at the University Hospital Düsseldorf from human patients in a sampling period between 2002 and 2020 (Fig. 3a, Supplementary Table 7). First, the strain collection of 113 isolates was characterised by multiplex polymerase chain reaction (PCR) to determine the distribution of *fpvA* type receptors and the overall prevalence of *pyoS2* and *imS2* genes within the strain library (Supplementary Fig. 7, Supplementary Table 7). Of the known ferripyoverdine siderophore receptors FpvAI, FpvAII, and FpvAIII, only FpvAI can be hijacked by PyoS2 for membrane translocation, rendering those strains with FpvAII- and FpvAIII inherently insensitive to PyoS2 treatment. At the same time, each strain encodes only one particular *fpvA*-type receptor^[38]^. Among the 113 tested strains, 15 strains (13.3%) were found to encode the *fpvAI* locus (Fig. 3b). Of these 15 strains belonging to the *fpvAI* siderovar, 8 strains (53.3%) additionally carried the *pyoS2*_*imS2* locus, rendering these strains capable of producing PyoS2 on their own and conferring resistance to wild-type PyoS2 by ImS2-mediated toxin neutralisation. In total, 83 strains (73.5%) within the strain library were shown to feature the *pyoS2_imS2* locus, making PyoS2 production a prevalent characteristic of the selected MDR microbes and underlining its role in microbial competition.

**Fig. 3.**
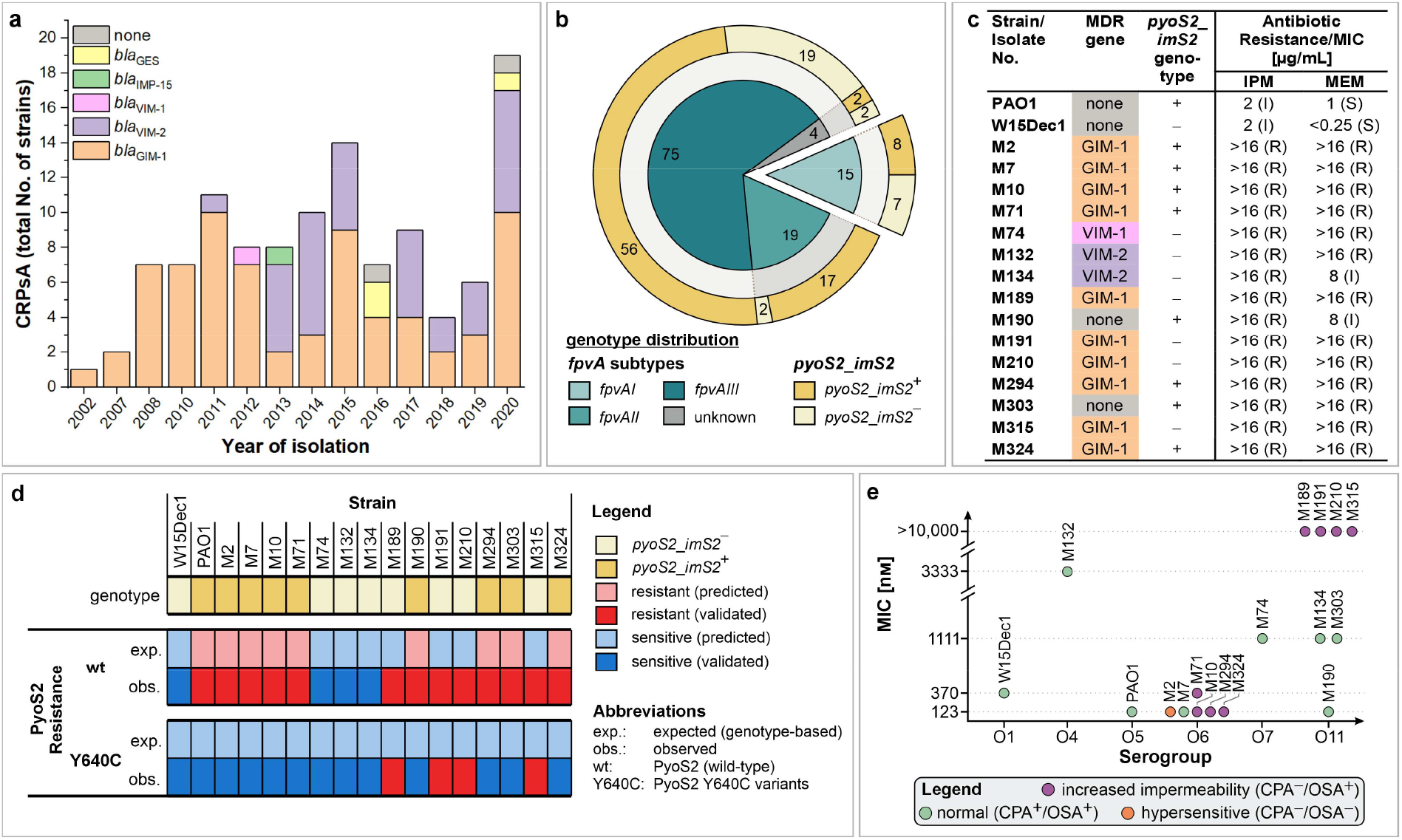
CRPsA screening and application of extended-spectrum PyoS2 antimicrobials on clinical isolates. **a**. Isolated CRPsA strains from the University Hospital in Düsseldorf (sampling period 2002−2020, n=113). **b**. Distribution of *fpvA* subtypes and *pyoS2*_*imS2* genotypes among the CRPsA strain collection (determined by multiplex PCR). **c**. List of *fpvAI*-positive strains (core strain collection) selected from the CRPsA collection. The *pyoS2*_*imS2* genotype is indicated, as well as identified carbapenemase genes and minimal inhibitory concentrations according to EUCAST guidelines against the carbapenem antibiotics imipenem (IPM) and meropenem (MEM). **d**. Expected and experimentally observed resistance profile of all strains from the core strain collection for PyoS2 (wild-type) and the PyoS2 Y640C variants with modifications at the IPE. **e**. Correlation between the serotype and the observed minimal inhibitory concentration of PyoS2 Y640C-AF488 against CRPsA strains (from agar spot assay).

Since only FpvAI-expressing strains are targetable with PyoS2, further evaluation of our extended-spectrum PyoS2 antimicrobials was focused on those 15 strains from the *fpvAI* siderovar (core strain collection, Fig. 3c), exhibiting accumulated carbapenem resistance caused by *bla*_GIM-1_, *bla*_VIM-1_, and *bla*_VIM-2_ metallo β-lactamase (carbapenemase) production, or by elevated membrane impermeability (only strains M190 and M303) (cf. Supplementary Table 8).

Phenotypic testing of the core strain collection of CRPsA isolates with wild-type PyoS2 led to the finding of only 3/7 *pyoS2_imS2*^−^ (42.9%) isolates being moderately sensitive to pyocin treatment (M74, M132, M134), even though these strains are not capable of producing the cognate immunity protein (Fig. 3d, Fig. 4). However, the other 4 strains (57.1%) sharing the *pyoS2_imS2*^−^ genotype exhibited an unexpectedly resistant phenotype (M189, M191, M210, M315). All remaining strains harbouring the *pyoS2_imS2*^+^ tandem met the expectation and showed ImS2-mediated resistance to PyoS2 treatment. As previously observed for the *P. aeruginosa* model strains, neither the cysteine mutants Q570C, K602C, and C-Cys, nor their corresponding AF488 conjugates were able to challenge the PyoS2 resistance of those strains carrying the *pyoS2_imS2*^+^ genotype (M2, M7, M10, M71, M190, M294, M303, M324). This PyoS2 resistance was effectively reversed for 8/8 strains (100%) by using the PyoS2 Y640C mutant and its AF conjugates, featuring the engineered IPE within PyoS2’s DNase domain.

**Fig. 4.**
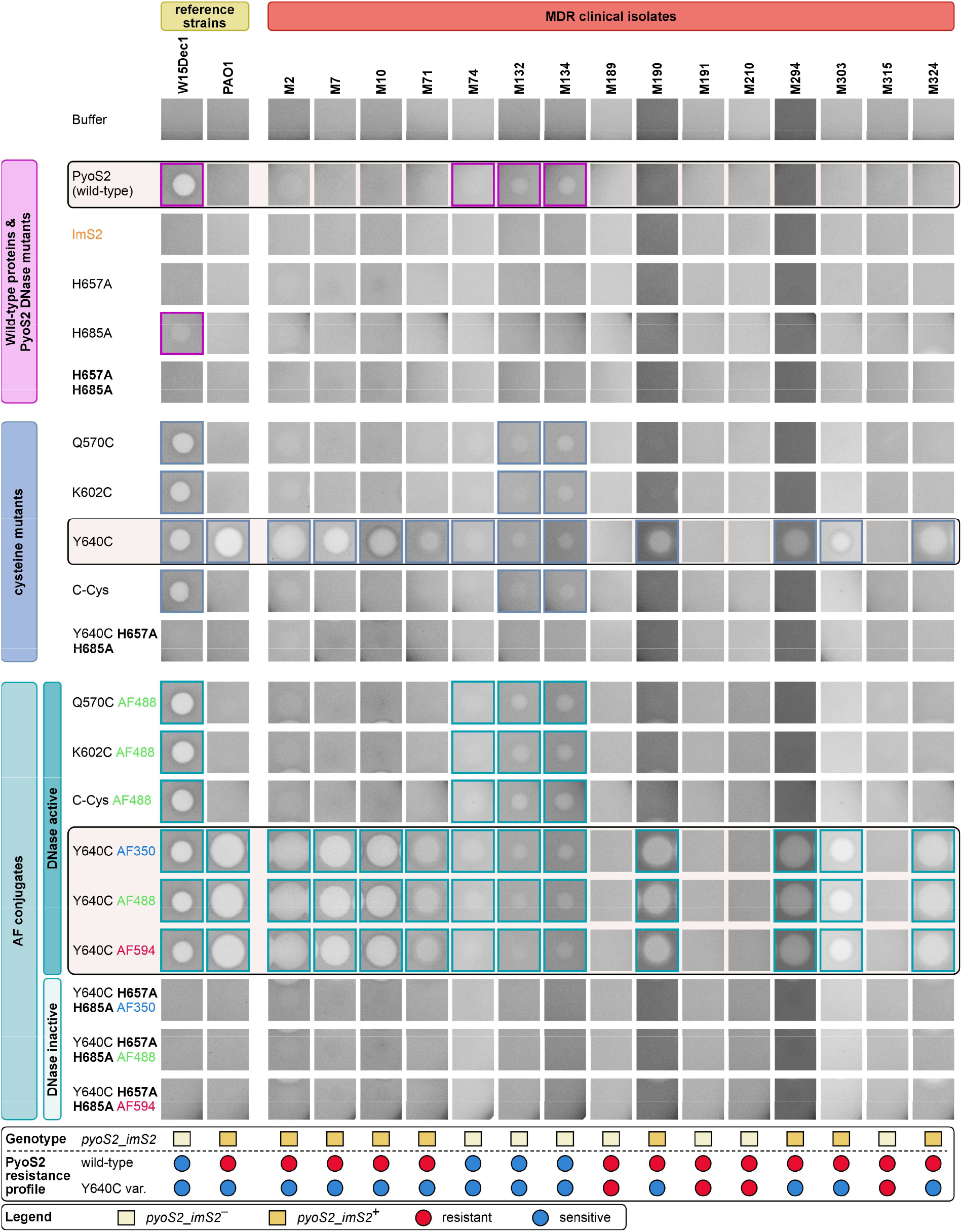
Killing activity of engineered PyoS2 derivatives against MDR *P. aeruginosa* isolates. Wild-type PyoS2, ImS2, and engineered PyoS2 mutants and conjugates were tested at 5 µM (5 µL) for killing activity against strain PAO1 (PyoS2 producer, PyoS2-resistant), strain W15Dec1 (PyoS2-sensitive), and *fpvAI*-positive MDR clinical isolates (M2−M324). Cells were immobilised in soft agar. Clear zones indicate inhibited bacterial growth.

While the extended-spectrum PyoS2 proteins were able to overrule ImS2-mediated PyoS2 resistance of all 8 *pyoS2_imS2*^+^ strains, thereby increasing the treatment success from 0% for PyoS2 to 100% for the PyoS2 Y640C variants, the 4 *pyoS2_imS2*^−^ strains M189, M191, M210, and M315 remained persistently resistant to the engineered toxin. Therefore, it was hypothesised that factors other than cognate immunity protein expression may cause this secondary PyoS2 resistance. Since PyoS2 translocation only involves the essential IM protease FtsH and the TonB1 complex at the inner membrane, and FpvAI and CPA at the outer membrane, whole-genome sequencing (WGS) was performed for strain W15Dec1 and all CRPsA strains from the core strain collection to search for genomic commonalities in the resistant isolates (cf. Fig. 3d). Using the tblastn algorithm, no recurring anomalies were found in the *ftsH, fpvA*, or *tonB1* loci that would explain the observed resistance patterns. Hence, further investigations were focused on cell-surface lipopolysaccharide biosynthesis, as the *P. aeruginosa* lipopolysaccharide antigens CPA and OSA (O-specific antigen) are known to have a pyocin-binding component^[21, 26, 39]^ or a cell-shielding^[40]^ character, respectively.

These bioinformatic analyses revealed recurring nonsense and frame-shift mutations in the *wzt* (strain M10), *wbpZ* (strains M294 and M324), and *gmd* loci (strains M2, M71, M189, M191, M210, and M315), all located within the CPA biosynthesis cluster of *P. aeruginosa* (Supplementary Fig. 8a). Wzt is the ATPase subunit of the Wzm/Wzt ABC transporter required for transport of the CPA from the cytoplasm to the outer leaflet of the IM (Supplementary Fig. 10)^[41]^. The cytoplasmic glycosyltransferase WbpZ initiates CPA polymerisation during CPA biosynthesis^[42-43]^. And the GDP-mannose dehydrogenase Gmd converts GDP-D-mannose to GDP-4-keto-6-deoxy-mannose^[44]^, which is an essential precursor of D-rhamnose. D-Rhamnose, in turn, is the direct precursor of the CPA D-rhamnose homopolymer and is required for the intracellular assembly of the CPA polysaccharide^[45]^ that functions as a secondary receptor on the cell surface for many S-type pyocins^[26, 39]^. Although all identified mutations render the respective CPA biosynthesis genes inactive pseudogenes and thus abolish CPA production, particularly strains carrying mutations within the *gmd* locus showed ambiguous effects on pyocin susceptibility. While the clinical isolates M189, M191, M210, and M315 were fully resistant to the extended-spectrum pyocins, the isolates M2 and M71 exhibited a sensitive phenotype.

Building on our earlier observation that PyoS2 Y640C variants produced larger zones of growth inhibition in strain PAO1 than in the PyoS2-sensitive isolate W15Dec1, we reasoned that the O serotype might influence the accessibility of CPA and the FpvAI receptor in the OM. Therefore, the serogroups of strains PAO1 and W15Dec1 were determined, based on the bacterial genomes decoded via WGS, by using the *Pseudomonas aeruginosa* serotyper 1.0 (PAst)^[46]^. While PAO1 was confirmed to belong to serotype O5^[47]^, W15Dec1 was identified to belong to the O1 serogroup, which substantiated the initial suspicion. Next, the minimal inhibitory concentrations (MICs) for wild-type PyoS2 and PyoS2 Y640C-AF488 were determined for all strains from the core strain collection (Supplementary Fig. 9), ranging from 123 nM to >10 µM (resistant). The subsequent *in silico* serotyping analysis uncovered the serogroups O1, O4, O5, O6, O7, and O11 within the core strain collection, of which O6 (6 strains) and O11 (7 strains) were best represented. By plotting the observed MICs for PyoS2 Y640C-AF488 treatment against the serotype (Fig. 3e), we were able to identify different clusters of ImS2-independent PyoS2 susceptibility. All strains with a persistent resistance to the extended-spectrum PyoS2 derivatives and defects in CPA biosynthesis through nonsense mutations in the *gmd* locus belonged to the O11 serotype (M189, M191, M210, and M315). In contrast, those strains with frame-shift mutations in the *gmd* locus belonging to the O6 serotype, namely M2 and M71, remained sensitive to the extended-spectrum pyocins.

Alongside mutations in the *gmd* locus, another frame-shift mutation was identified in the *wbpL* locus of strain M2 (Supplementary Fig. 8b). WbpL is a cytoplasmic glycosyltransferase, which catalyses the initial step in both CPA and OSA assembly (Supplementary Fig. 10a)^[42]^. Consequently, strain M2 is not only deficient for the CPA antigen but also for the OSA antigen, which causes this strain to become hypersensitive to treatment with the extended-spectrum pyocins (cf. Supplementary Fig. 8a).

In summary, the data suggest that O-specific antigens play a significant role in modulating the accessibility of the OM for environmental toxins such as S-type pyocins. In the presence of the CPA lipopolysaccharide, this effect fades into the background, as CPA binding enhances the affinity of PyoS2 to the target cell’s membrane. However, in the absence of CPA, the O11 serotype apparently provides a denser packing of the OM and thereby effectively shields the FpvAI receptor from PyoS2 binding. The O6 serotype, however, does not offer such efficient shielding, thereby enabling efficient PyoS2 binding and killing, even in the absence of the CPA lipopolysaccharide. Similar effects were previously described for phage-like R-type pyocins^[40]^, although R-type pyocins primarily bind to sugar motifs found in the LPS core region rather than the D-rhamnose-containing CPA lipopolysaccharide.

### HNH DNase bacteriocins and their immunity proteins employ a conserved topology

In this case study, we clearly demonstrated that the antimicrobial spectrum of PyoS2 can be expanded by preventing its interaction with the cognate immunity protein ImS2. Since the formation of toxin-antitoxin pairs is not unique to PyoS2 and widespread among Gram-negative bacteriocins, we hypothesised that other bacteriocin/immunity protein pairs with homologous HNH DNase effectors can likewise be manipulated to unleash effector cytotoxicity insensitive to immunity protein binding and thereby broaden their antimicrobial activity.

The sequence comparison of several HNH DNase domains from colicins, klebicins, and pyocins earlier revealed that the sequence conservation at the immunity protein exosite – comprising helix V, loop 4, 3_10_ helix, loop 5, β-strand 3, and the beginning of loop 6 (cf. Fig. 1b) – is overall lower than for the other fractions of the DNase domain. This observation can be explained by the fact that the remainder of the residues is required to fold into the conserved HNH nuclease fold, while those residues at the IPE are utilised to mediate selective binding to sequence-wise unique cognate immunity proteins. Despite the high degree of sequence variation and low sequence homology (for selected homologies see Fig. 5g), DNase and Im association in ultra-high-affinity complexes resorts to a conserved topology, as reflected by the very high structural congruency of DNase and ImS2 structures (RMSD <1 Å). A comparative analysis of crystal structures of four HNH nuclease bacteriocins, namely PyoS2, ColE2, ColE7, and PyoAP41, in complex with their cognate immunity proteins, clearly demonstrates that the specific residues and their interacting partners may be different, but at the same time, the overall architecture and topology of the protein-protein interface are highly conserved (Fig. 5a−d). The central residue – equivalent to Y640 in PyoS2 − is laterally stabilised by two DNase side chains (−8 and +11 positions from the central DNase residue) and axially interacts with the side chain of a residue directly interacting with the DNA substrate (+12 positions from the central DNase residue) (Fig. 5e, top). Likewise, the relative position of Im residues interacting with the DNase is conserved in the primary sequence of the immunity proteins. Starting from a conserved His-Pro motif, four residues were identified to interact with the central DNase residue. A conserved aromatic residue (+8 positions from the conserved His residue) supports the central DNase residue from below, while residues in the relative positions −13, −12, and −9, all located in the energy binding hotspot helix II of the immunity protein, directly face the central DNase residue (Fig. 5e, bottom). Based on this conserved complex topology, we developed an illustrative model highlighting the geometry of the protein interface around the central DNase residue (Fig. 5f).

**Fig. 5.**
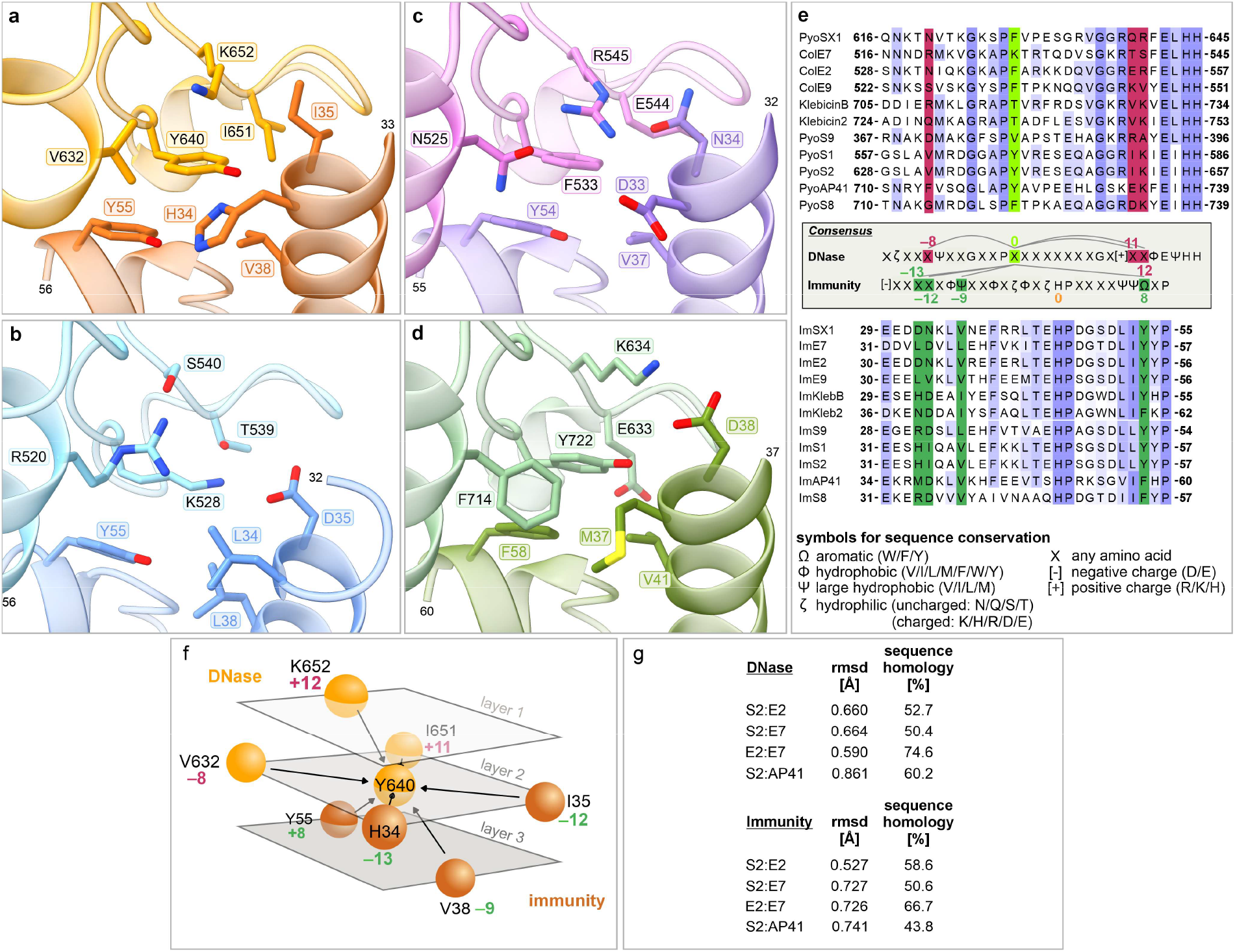
Structural analysis of IPEs from representative HNH DNase bacteriocins, highlighting the conserved topology at the protein interface. **a**. IPE excerpt from the PyoS2/ImS2 crystal structure (4QKO). **b**. IPE excerpt from the ColE2/ImE2 crystal structure (3U43). **c**. IPE excerpt from the ColE7/ImE7 crystal structure (7CEI). **d**. IPE excerpt from the PyoAP41/ImAP41 crystal structure (4UHP). **e**. Multiple sequence alignment of numerous HNH DNase IPEs and immunity proteins. The ‘central’ DNase residue (equivalent to Y640 in PyoS2) is highlighted in bright green and flanking residues within the DNase are marked in purple. Interacting residues within the immunity protein are highlighted in dark green. **f**. Three-layered model showcasing the conserved topology of side chain interactions around the ‘central’ residue. **g**. Comparison of HNH DNase domains and cognate immunity protein sequence homology and structural conservation of the protein backbones from **a**−**d**, revealing poor sequence conservation but exceptionally high structural congruency.

Based on this model, it can be assumed that the principle of weakened PyoS2/ImS2 affinity uncovered in this study might be transferable to other HNH DNase bacteriocins as well. As the loop 4 residue Y640 from the IPE of PyoS2 was identified as a pivotal residue essential for mediating tight binding to ImS2, the replacement of equivalent residues within other HNH DNase bacteriocins in this particular central position could potentially engender a general means of antimicrobial spectrum expansion and thereby curtail the practical relevance of cognate immunity proteins and immunity protein-mediated bacteriocin resistance.

## Discussion

In this study, we used the HNH DNase bacteriocin pyocin S2 from *P. aeruginosa* to assess its potential as an antimicrobial agent against pathogenic *P. aeruginosa* strains, including highly relevant carbapenem-resistant clinical isolates. By introducing point mutations at PyoS2’s IPE and blocking the association of the ultra-high-affinity complex between PyoS2 and its cognate immunity protein ImS2 through chemical functionalisation via maleimide chemistry, we generated the first known extended-spectrum pyocins that are not based on chimeric architecture and domain swapping^[48]^. In this regard, a single Y640C mutation was sufficient to overrule ImS2-mediated PyoS2 resistance in all tested *P. aeruginosa* model strains and CRPsA isolates belonging to the *fpvAI* siderovar, and the attachment of non-proteinogenic fluorophore labels even more enhanced the effect by rescuing DNase activity. While the generation of chimeric bacteriocins with shuffled receptor-targeting and cytotoxic domains from different bacteriocins and various organisms^[11, 48]^ drives the evolutionary pressure for challenged microbes to acquire the corresponding immunity proteins from environmental sources, the thiol-maleimide click chemistry and the fluorophores used in this study are fully artificial and easily substitutable, respectively. This circumstance can be taken advantage of to alter the protein-protein interface and reduce the risk of emerging escape mutants relying on spontaneous counter-mutations in the cognate immunity protein. Moreover, by shuffling receptor binding domains that target more prevalent OM receptors (e.g. Hur, CrtA, FpvA; Supplementary Fig. 10b) and combining them with the extended-spectrum DNase domain of PyoS2, novel chimeric pyocins can be generated that potentially address an even larger fraction of *P. aeruginosa* strains. Additionally, the successful cellular import of PyoS2-AlphaFluor conjugates could be used as a starting point for developing bacteriocins and their conserved TonB-dependent import mechanisms as novel drug vectors that circumvent the prevalent drug efflux mechanisms leading to multidrug resistance.

Another mechanism of escape from the antimicrobial activity of PyoS2, besides the expression of ImS2, was found in the abolishment of CPA biosynthesis, removing the secondary PyoS2 receptor CPA from the OM. The virulence factor CPA of *P. aeruginosa* is required for the maturation of robust biofilms^[49]^ and functions as a receptor for conjugative type IV pili^[50]^, which are required for the horizontal gene transfer of the PAPI-1 pathogenicity island to recipient strains. However, the loss of the CPA lipopolysaccharide antigen through spontaneous mutations in the corresponding gene cluster can increase the fitness of *P. aeruginosa* strains by evading the cytotoxic influence of S-type pyocins in competitive ecological niches. Alterations of LPS are generally a widely used strategy in the bacterial kingdom to increase prokaryotic fitness under certain stress factors. For the human pathogen *P. aeruginosa*, mutations in loci related to LPS biosynthesis have been previously shown to enhance resistance to β-lactams^[51-52]^ and to increase fitness in co-cultures with *Staphylococcus aureus*^[52]^. Therefore, a dysfunctional CPA biosynthesis cluster in clinical isolates may not only help reduce susceptibility to S-type pyocins but also increase the tolerance to common antibiotics and facilitate the co-evolution with other human pathogens in immunocompromised patients.

Moreover, our study highlights the serotype-dependence of bacteriocin killing, which has been previously observed in *E. coli* with colicins^[53]^ and in *P. aeruginosa* with R-type pyocins^[40]^. However, colicins do not require the secondary CPA receptor to be present, as this antigen is a unique trait of *Pseudomonads*^[43]^. Also, R-type pyocins are phage-like tailocins for which no immunity proteins are required. Thus, the serotype-dependent component of S-type pyocin killing and insensitivity in *P. aeruginosa* was so far concealed by immunity protein-mediated resistance and CPA-deficient phenotypes. The novel PyoS2 Y640C derivatives with defects in ImS2 binding bring this matter to light for the first time, accentuating once more the requirement of understanding the biosynthesis, morphology, and the cell surface-shielding effect of O-specific antigens in bacterial human pathogens. Moreover, these presented results call for the development of therapeutic LPS-degrading glycosidases or lyases, which might, for example, find application in intranasal administration^[54-55]^ or salve^[56]^ co-formulations with pyocins. This combinatorial treatment could facilitate the bacteriocin uptake into target cells with dense OSA packing (e.g. serotype O11) in multidose treatments by remodelling the exposed lipopolysaccharides – even in the absence of the secondary CPA receptor. Enzymes with OSA-degrading activity can be found in bacteriophages, which frequently recognise OSA for initial cell adhesion and can dismantle the OSA for facilitated infectivity. One such enzyme (LKA1gp49) has been found recently in the tail spike of the *P. aeruginosa*-specific phage LKA1, degrading the O5-specific polysaccharide through lyase activity^[57]^.

While this case study was solely focused on the best-studied S-type pyocin S2, the presented findings challenge the general concept of immunity protein-mediated bacteriocin resistance and have, therefore, broader implications for the general application of Gram-negative bacteriocins as antimicrobial agents in the current antimicrobial resistance crisis. Notably, the exceptionally high structural conservation of HNH DNase domains suggests transferability of the uncovered mechanisms of antimicrobial spectrum expansion to other HNH bacteriocins and Gram-negative organisms. Finally, we have presented a precedent for overcoming the immunity protein-mediated limitations of Gram-negative bacteriocins, paving the way for their comprehensive long-term application for the treatment of nosocomial infections with drug-resistant, difficult-to-treat human pathogens.

## Acknowledgements

T.M.W. was supported with a doctoral scholarship by the Jürgen Manchot Foundation. The authors thank Prof. em. Pierre Cornelis and Sandra Matthijs for providing the PyoS2-sensitive environmental isolate of *P. aeruginosa* W15Dec1. The authors also wish to thank Nadiia Pozhydaieva for her valuable contributions to the processing and annotation of the bacterial genomes and her substantial feedback regarding this manuscript. Additionally, credit is ascribed to Jan-Hendrik Illies and Maria Emilia Iglesias Moncayo for their diligent proofreading. Lastly, the Heinrich Heine University Düsseldorf and Forschungszentrum Jülich are acknowledged for their ongoing support.

## Author contributions

**T.M.W**. – project idea, project conceptualisation, and project administration, funding acquisition, experimental work (except specified below), data analysis, visualisation, conceptualisation and writing of the manuscript. **R.G**. – experimental work (isolation of MDR *P. aeruginosa* strains and characterisation of carbapenem resistance by RT-PCR and antibiotic susceptibility testing. **C.R.M**. – project funding. **J.P**. – project funding and supervision. All authors reviewed and approved the final paper.

## Declaration

The utilisation of clinical isolates of MDR strains of *Pseudomonas aeruginosa* from hospitalised human patients with the study number **2021-1324** (*Microbiological evaluation of pyocin S2 conjugates as substitutes for antibiotics and the treatment of multidrug-resistant Gram-negative bacteria*) was reviewed and approved by the ethics commission of the Faculty of Medicine from the Heinrich Heine University Düsseldorf.

## Additional information

**Supplementary information**is available for this paper. The authors have cited additional references within the supplementary information^[58-91]^.

