## Supplementary Information for "Hoisted by Their Own Petard: Defeating Immunity Protein–Mediated Bacteriocin Resistance"

### Supplementary data

**Supplementary Table 1.** Predicted and experimentally validated S-type pyocins, their OM receptors, and their pathway of translocation.

| Pyocin | Killing Activity | OM Receptor | Translocation Pathway | Reference |
|---|---|---|---|---|
| PyoS1 | DNase (HNH) | unknown | TonB1-dependent | [1] |
| PyoS2 | DNase (HNH) | FpvAI | TonB1-dependent | [1-3] |
| PyoS3 | DNase | FpvAII | TonB1-dependent | [4-5] |
| PyoS4 | tRNase | FpvAI | TonB1-dependent | [6-7] |
| PyoS5 | pore-forming | FptA | TonB1-dependent | [8-10] |
| PyoS6 | rRNase | unknown | unknown | [11] |
| PyoS7* | rRNase | FpvAI† | TonB1-dependent | [12] |
| PyoS8 | DNase (HNH) | unknown | unknown | [13] |
| PyoS9** | DNase (HNH) | unknown | unknown | [12] |
| PyoS10*** | DNase | unknown | unknown | [12] |
| PyoAP41 | DNase (HNH) | unknown | Tol-dependent | [14] |
| PyoG | nuclease | Hur | TonB1-dependent | [15] |
| PyoSX1 | DNase (HNH) | CrtA | TonB1-dependent | [16] |
| PyoSX2 | inhibition of protein synthesis | CrtA | TonB1-dependent | [16] |
| PyoS11/PyoSD2 | tRNase | FpvAI | TonB1-dependent | [17] |
| PyoS12/PyoSD3 | tRNase | FpvAII‡ | TonB1-dependent | [17] |
| PyoS13/PyoSD1 | tRNase | unknown | TonB1-dependent | [17] |

* Predicted from *P. aeruginosa* BWHPSA018; not experimentally validated. † First 464 amino acids are identical to PyoS2 from *P. aeruginosa* PAO1, why FpvAI must be the OM receptor of PyoS7. ** Predicted from *P. aeruginosa* BL04. *** Predicted from *P. aeruginosa* PABL056. ‡ High sequence homology between the N-terminal domains of PyoSD3 and PyoS3 suggests translocation via the FpvAII siderophore receptor. However, experimental proof is missing.

**Supplementary Table 2**. Polar interaction of amino acid residues at the interface between PyoS2 and ImS2. Interactions and distances have been collected from the PDB-REDO structure 4QKO using PDBePISA[18]. Distances were measured between heteroatoms, omitting interjacent hydrogen atoms. Abbreviations: BB – backbone; SC – side chain.

| PyoS2 | | | ImS2 | | | |
|---|---|---|---|---|---|---|
| **Hydrogen bonds** | | | | | | |
| **Residue** | **Position** | **Atom name** | **Residue** | **Position** | **Atom name** | **Distance [Å]** |
| N626 | SC | ND2 | Y21 | SC | OH | 3.78 |
| | SC | ND2 | L54 | BB | O | 2.89 |
| | SC | ND2 | D63 | SC | OD1 | 3.22 |
| | SC | ND2 | D63 | SC | OD2 | 3.22 |
| A631 | BB | O | N23 | SC | ND2 | 3.13 |
| D635 | SC | OD2 | N23 | SC | OD1 | 3.31 |
| | SC | OD2 | K25 | SC | NZ | 2.75 |
| Y640 | SC | OH | H34 | SC | ND1 | 2.75 |
| | BB | O | Y56 | SC | OH | 2.64 |
| E643 | SC | OE1 | K42 | SC | NZ | 3.26 |
| | SC | OE2 | K42 | SC | NZ | 2.96 |
| | SC | OE2 | S51 | BB | N | 2.88 |
| | SC | OE2 | S51 | SC | OG | 3.20 |
| | BB | N | D52 | SC | OD1 | 2.78 |
| Q646 | SC | NE2 | S51 | SC | OG | 2.82 |
| | SC | OE1 | K42 | SC | NZ | 3.34 |
| K652 | SC | NZ | E31 | SC | OE2 | 2.66 |
| **Salt bridges** | | | | | | |
| **Residue** | **Position** | **Atom name** | **Residue** | **Position** | **Atom name** | **Distance [Å]** |
| R605 | SC | NH2 | E32 | SC | OE1 | 3.51 |
| | SC | NH1 | E32 | SC | OE2 | 3.98 |
| | SC | NH2 | E32 | SC | OE2 | 2.98 |
| R608 | SC | NH1 | E31 | SC | OE1 | 2.92 |
| | SC | NH2 | E31 | SC | OE1 | 3.62 |
| | SC | NH1 | E31 | SC | OE2 | 3.47 |
| | SC | NH1 | E31 | SC | OE1 | 2.65 |
| D635 | SC | OD2 | K25 | SC | NZ | 2.75 |
| E643 | SC | OE1 | K42 | SC | NZ | 3.26 |
| | SC | OE2 | K42 | SC | NZ | 2.96 |
| K652 | SC | NZ | E31 | SC | OE2 | 2.66 |

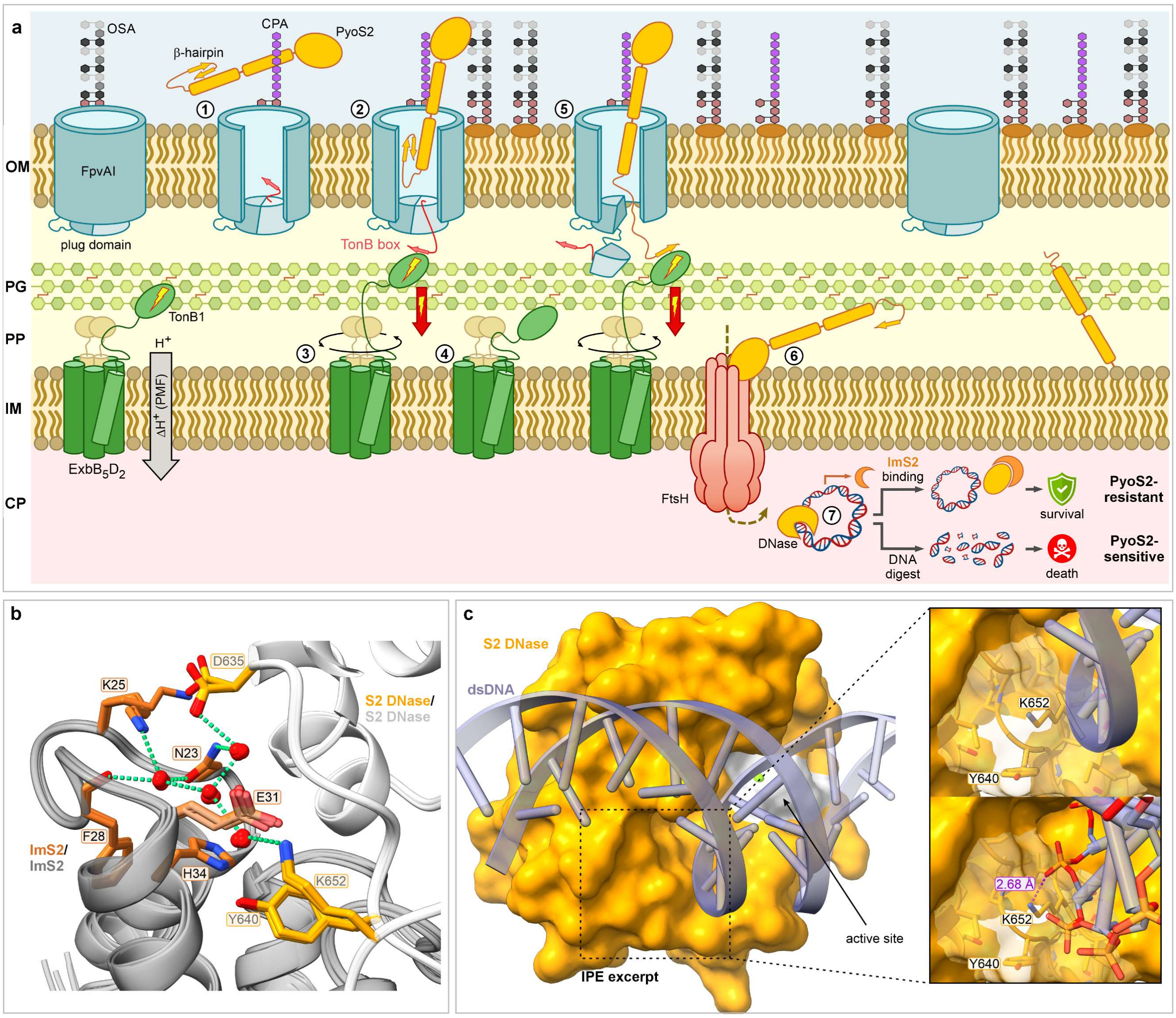


**Supplementary Fig. 1.** Adapted import mechanism and mode of action of PyoS2. **a**. **1**. The flexible N-terminus of S-type pyocins S2 is folded in a β-hairpin and docked to the bundle of helix I and V of the bacterial toxin. The TonB-dependent transporter (TBDT) FpvAI is plugged on the periplasmic side through a plug domain that contains a force-resistant (blue) and force-labile (light blue) subdomain. The TonB box within the plug domain is buried in the lumen of the TBDT. **2**. Upon binding of the pyocin to the primary TBDT and the secondary CPA receptor, a conformational change in the plug domain is induced, uncovering the TonB box. **3**. The periplasmic portion of TonB1 from the Ton machinery is recruited and interacts with the TonB box of the TBDT through β-augmentation. Movement of the Ton motor removes the force-labile subdomain of the TBDT plug. **4**. Discharged TonB1 is recharged for subsequent motor movement. **5**. The highly flexible N-terminus of the S-type pyocin enters the periplasm and its TonB box aligns with the periplasmic portion of TonB1 through β-augmentation. Movement of the Ton motor drags the pyocin through the pore of the TBDT for translocation of the toxin into the target cell's periplasm. **6**. The inner membrane AAA$^{+}$ protease FtsH cleaves the C-terminal cytotoxic domain from the helical receptor binding and translocation domain in an unknown mechanism. Only the nuclease domain is guided into the cytoplasm. **7**. The C-terminal HNH DNase domain displays cytotoxic activity by unspecific hydrolytic cleavage of intracellular DNA. The breakdown of nucleic acid metabolism causes highly potent cell killing. However, strains producing the cognate immunity protein ImS2 neutralise the toxin by ultra-high-affinity binding without being harmed. **b**. Display of interstitial waters at the PyoS2/ImS2 interface, conserved within all four asymmetric units of the crystal unit cell. **c**. AlphaFold model of the PyoS2 DNase bound to a $[AT]_9$ dsDNA substrate showing the interaction of K652 with the DNA backbone. The DNase active site is highlighted in white. The computed AlphaFold model aligns with the crystallised ColE7/DNA complex (PDB 1ZNS) with an RMSD of 0.641 Å (not shown). Abbreviations: CP – cytoplasm; CPA – common polysaccharide antigen; IM – inner membrane; OM – outer membrane; OSA – O-specific antigen; PG – peptidoglycan; PP – periplasm.

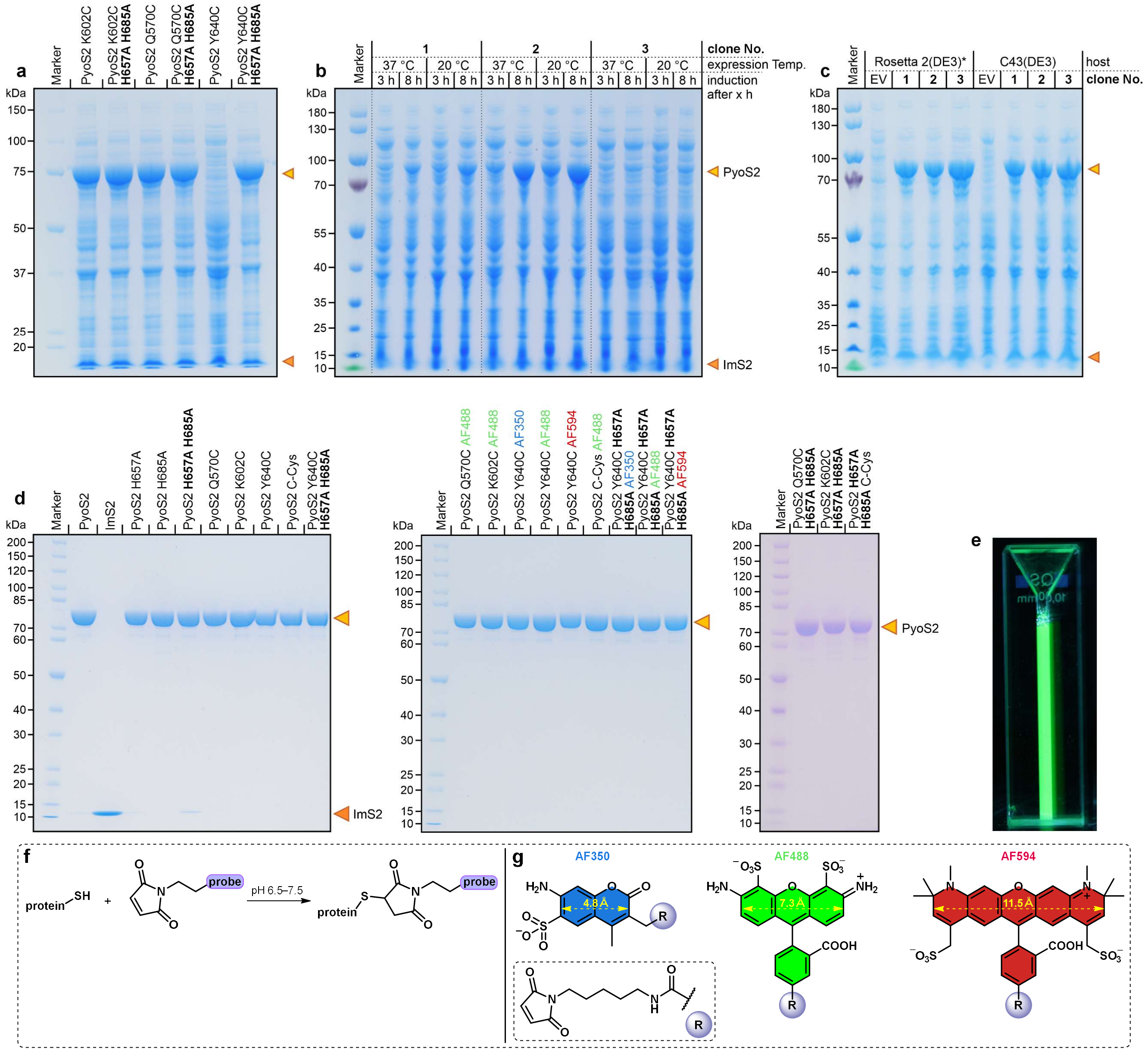


**Supplementary Fig. 2.** SDS-PAGE gels of PyoS2 expression and purified PyoS2 proteins, as well as thiol-maleimide labelling chemistry. Proteins were separated under denaturing conditions on 4–12% bis-tris PAGE gels at a loading of $OD_{600}$ 0.1/well (whole cells) or 2 µg (purified proteins). **a**. Heterologous expression of various PyoS2 cysteine mutants in *E. coli* BL21(DE3). **b**. Inhomogeneous expression of PyoS2 Y640C from three individual clones of BL21(DE3) with varying time points of induction (3 h vs. 8 h) and expression temperatures (20 °C vs. 37 °C). **c**. Reliable expression of PyoS2 Y640C from *E. coli* Rosetta 2(DE3) pLysS or *E. coli* C43(DE3)[1]. **d**. SDS-PAGE gels of purified PyoS2, ImS2, PyoS2 mutants and PyoS2-AlphaFluor conjugates. **e**. Green fluorescence of PyoS2 Y640C AF488 after irradiation with UV light (365 nm). Protein sizes: ImS2 – 11 kDa; PyoS2 – 74 kDa. *Rosetta 2(DE3) pLysS. **f**. Michael addition of a thiol-containing protein to a maleimide moiety with an arbitrary probe, yielding covalent protein-maleimide conjugates. **g**. Structure of AlphaFluor 350 (AF350), AlphaFluor 488 (AF488), and AlphaFluor 594 (AF594) $C_5$ maleimides. The ring colour corresponds to the colour at the absorbance maximum. Diameters were measured with Chem3D and do not include ring substituents.

[1] JM109(DE3) did not even grow in liquid culture, Rosetta 2(DE3) and C43(DE3) both were able to produce the protein upon induction. However, since strain C43(DE3) exhibited significantly improved growth in contrast to Rosetta 2(DE3), as reflected by the 2.6-times higher final optical density ($OD_{600}$ 15.6 ± 0.3 vs. 6.1 ± 0.2), strain C43(DE3) with its intrinsically high tolerance for toxic proteins was employed for the production of PyoS2 Y640C.

**Supplementary Table 3.** ESI-MS data for ImS2, PyoS2, PyoS2 mutants, and PyoS2-AF conjugates. Abbreviation: SD – standard deviation.

| Protein | Average Mass*† [calc.] | Mass [detect.] | SD |
|---|---|---|---|
| ImS2-$His_6$ | 11104.47 | 11104.60 | - |
| PyoS2 | 73722.52 | 73722.90 | 0.88 |
| PyoS2 H657A | 73656.46 | 73658.58 | 2.33 |
| PyoS2 H685A | 73656.46 | 73655.83 | 4.42 |
| PyoS2 **H657A H685A** | 73590.40 | 73590.98 | 0.76 |
| PyoS2 Q570C | 73697.55 | 73697.56 | 2.06 |
| PyoS2 Q570C AF488 | 74394.53 | 74391.22 | 2.17 |
| PyoS2 K602C | 73697.61 | 73694.19 | 6.19 |
| PyoS2 K602C AF488 | 74394.61 | 74394.15 | 3.13 |
| PyoS2 Y640C | 73662.61 | 73662.87 | 2.15 |
| PyoS2 Y640C AF350 | 74138.09 | 74136.08 | 0.11 |
| PyoS2 Y640C AF488 | 74359.60 | 74359.12 | 2.63 |
| PyoS2 Y640C AF594 | 74548.58 | 74551.04 | 1.57 |
| PyoS2 Y640C **H657A H685A** | 73530.48 | 73530.92 | 1.67 |
| PyoS2 Y640C **H657A H685A** AF350 | 74006.96 | 74005.86 | 1.34 |
| PyoS2 Y640C **H657A H685A** AF488 | 74228.14 | 74227.03 | 1.38 |
| PyoS2 Y640C **H657A H685A** AF594 | 74416.45 | 74415.45 | 6.04 |
| PyoS2 C-Cys | 73825.78 | 73825.75 | 1.51 |
| PyoS2 C-Cys AF488 | 74519.79 | 74519.85 | 2.49 |

* All proteins and masses were calculated and detected with the initial formyl-methionine missing.

† For PyoS2-AlphaFluor conjugates, the average mass was calculated from the PyoS2 molecular weight without initial methionine and masses of the commercial AlphaFluor350 (578.68 g/mol triethylammonium counter ion), AlphaFluor488 (720.66 g/mol, sodium counter ion), and AlphaFluor594 (908.97 g/mol, sodium counter ion) $C_5$ maleimides. A mass of 23 was subtracted from the indicated molecular weight for sodium as the counter ion, and a mass of 102.20 for the triethylammonium counter ion.

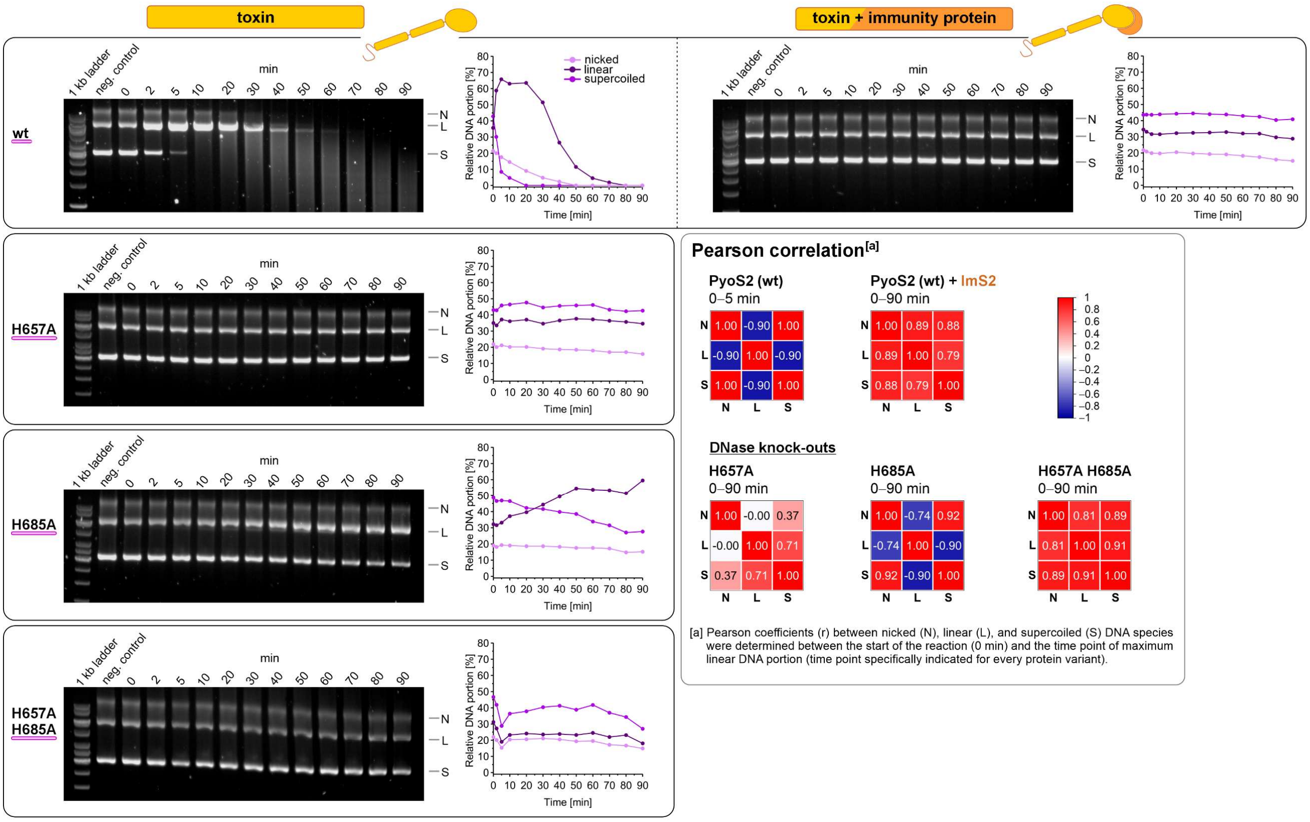


**Supplementary Fig. 3.** DNase kinetics of PyoS2, its neutralisation by ImS2, and kinetics of DNase active site mutants. DNA digest of pET21a(+) plasmid DNA with 5 µM PyoS2 or a mixture of 5 µM PyoS2 and 5.25 µM ImS2 was monitored over 90 min and analysed by agarose gel electrophoresis. Graphs were generated by densitometric analyses and display the relative portion of DNA (nicked, linear, supercoiled) in relation to incubation time. Pearson coefficients were determined from 0 min to the time point where the highest linear DNA portion was observed to distinguish assay-related coincident decrease of all DNA species (no DNase activity) from real DNase activity (particularly negative correlation between supercoiled and linear DNA). Fit parameters are summarised in Supplementary Table 4, 5, and 6.

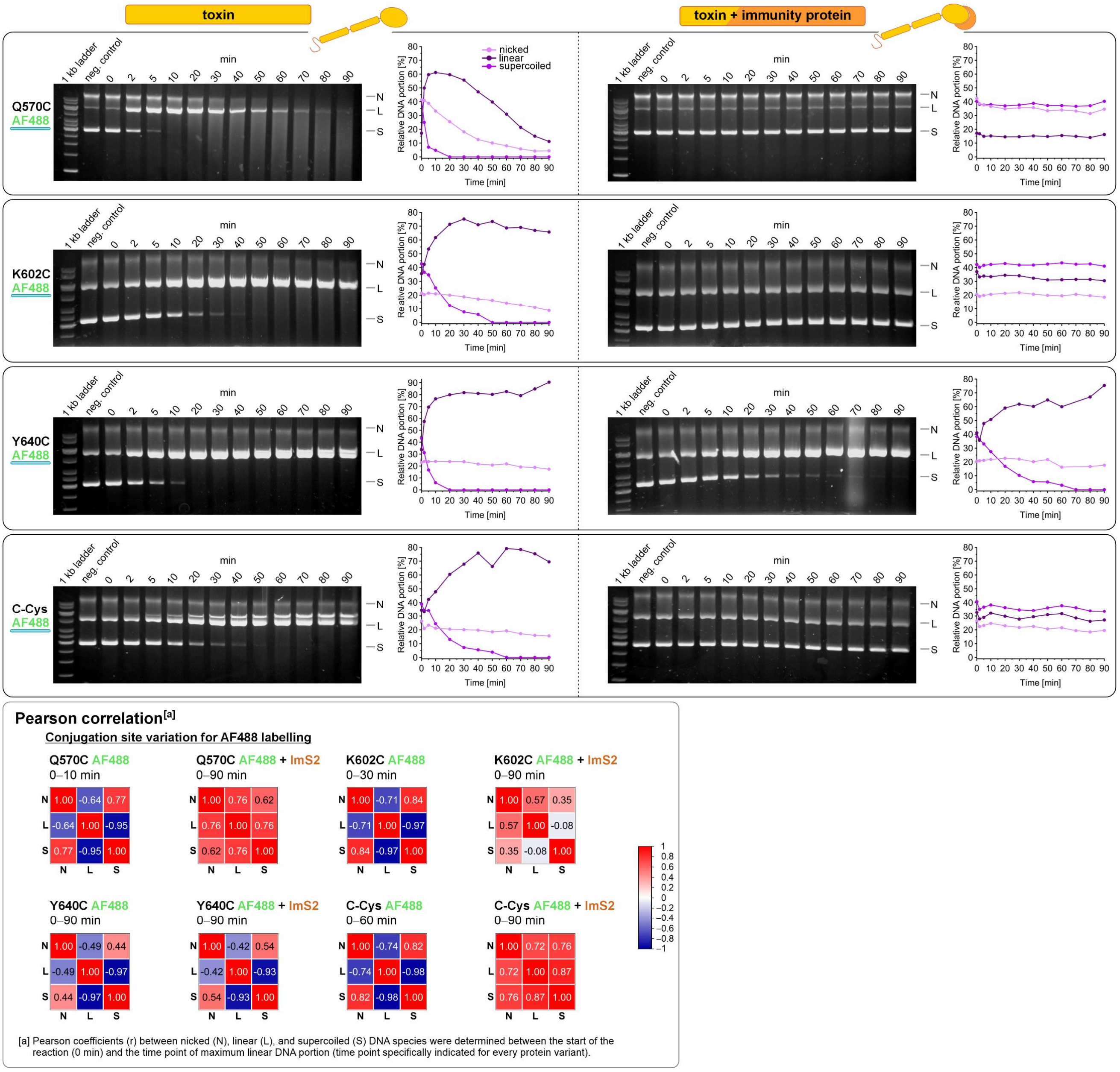


**Supplementary Fig. 3 (continued).** DNase kinetics of PyoS2-AF488 conjugates and their neutralisation by ImS2. DNA digest of pET21a(+) plasmid DNA with 5 µM PyoS2 or a mixture of 5 µM PyoS2 and 5.25 µM ImS2 was monitored over 90 min and analysed by agarose gel electrophoresis. Graphs were generated by densitometric analyses and display the relative portion of DNA (nicked, linear, supercoiled) in relation to incubation time. Pearson coefficients were determined from 0 min to the time point where the highest linear DNA portion was observed to distinguish assay-related coincident decrease of all DNA species (no DNase activity) from real DNase activity (particularly negative correlation between supercoiled and linear DNA). Fit parameters are summarised in Supplementary Table 4 and 5.

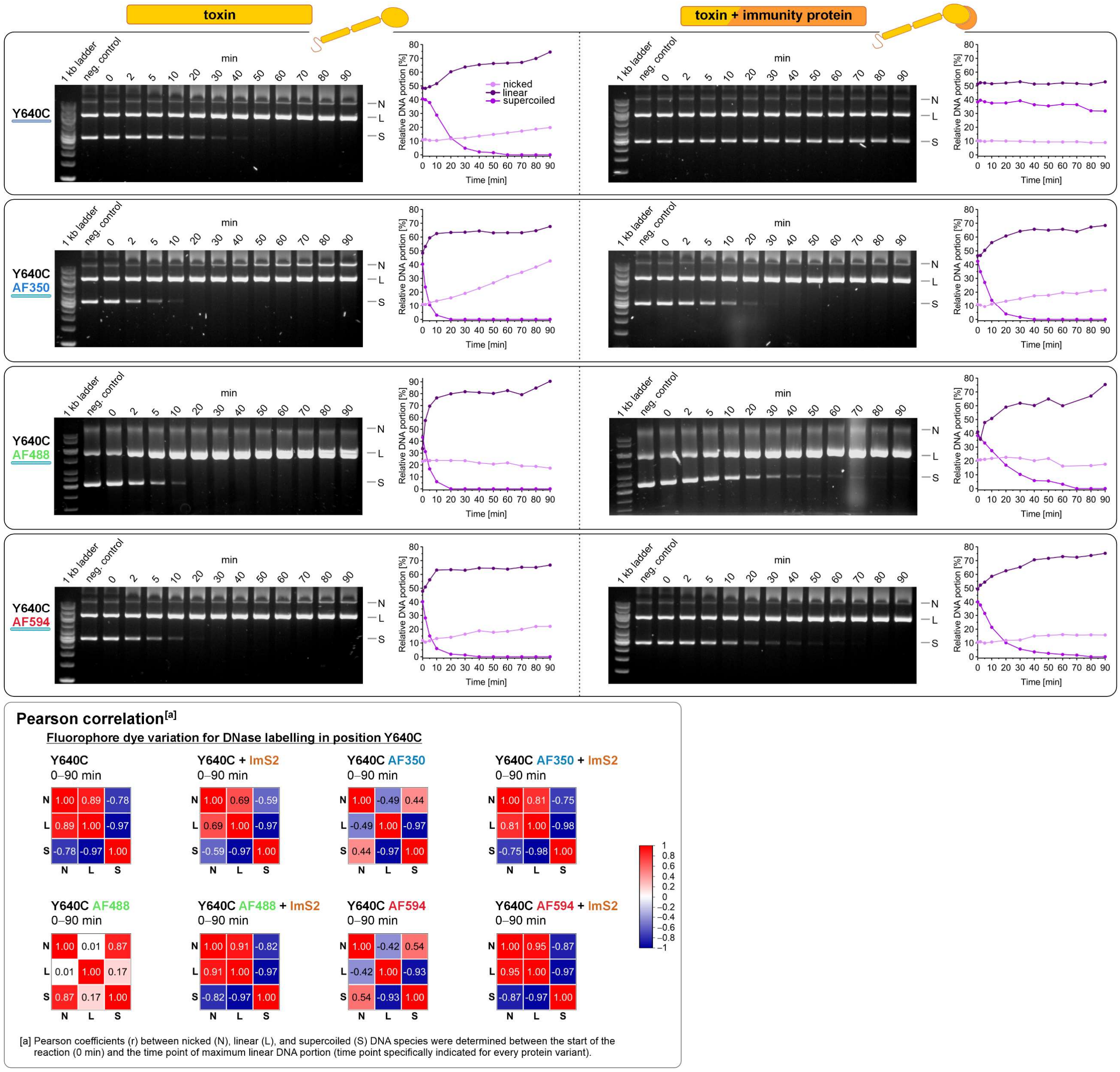


**Supplementary Fig. 3 (continued).** DNase kinetics of PyoS2 Y640C, its AF dye conjugates, and their neutralisation by ImS2. DNA digest of pET21a(+) plasmid DNA with 5 µM PyoS2 or a mixture of 5 µM PyoS2 and 5.25 µM ImS2 was monitored over 90 min and analysed by agarose gel electrophoresis. Graphs were generated by densitometric analyses and display the relative portion of DNA (nicked, linear, supercoiled) in relation to incubation time. Pearson coefficients were determined from 0 min to the time point where the highest linear DNA portion was observed to distinguish assay-related coincident decrease of all DNA species (no DNase activity) from real DNase activity (particularly negative correlation between supercoiled and linear DNA). Fit parameters are summarised in Supplementary Table 4 and 6.

**Stretched exponential decay**

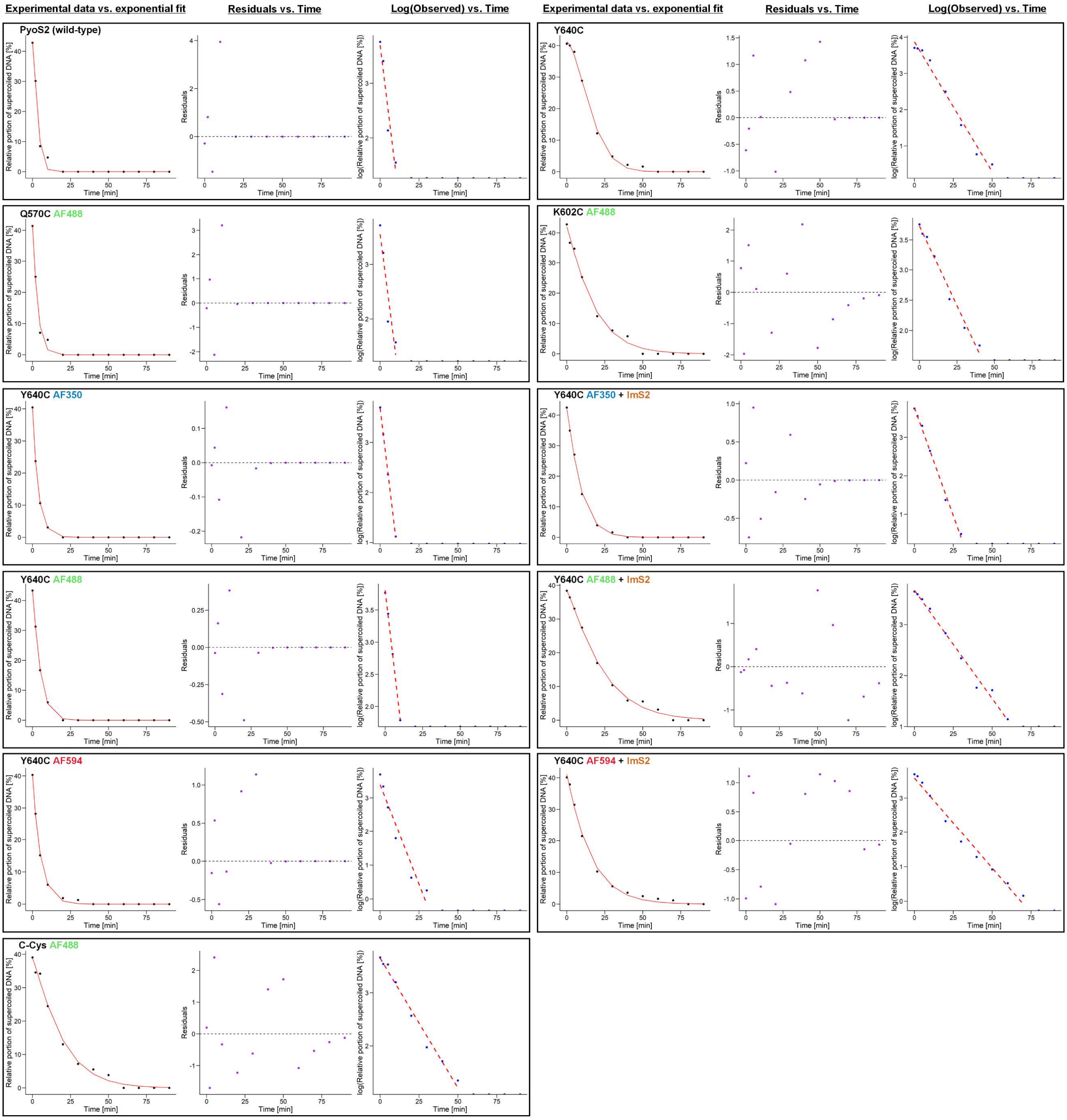


**Supplementary Fig. 4.** Fitting of supercoiled DNA depletion to an exponential decay function. The relative supercoiled DNA portion of PyoS2 mutants and conjugates with high DNase activity (data taken from Supplementary Fig. 3) are fitted to the Kohlrausch-Williams-Watts function (stretched exponential decay). Shown are three individual plots: i) experimental data vs. fit, ii) residuals vs. time, and iii) log(observed) vs. time. Fit parameters and the estimated half-life of supercoiled DNA are summarised in Supplementary Table 4.

#### Permutation analysis

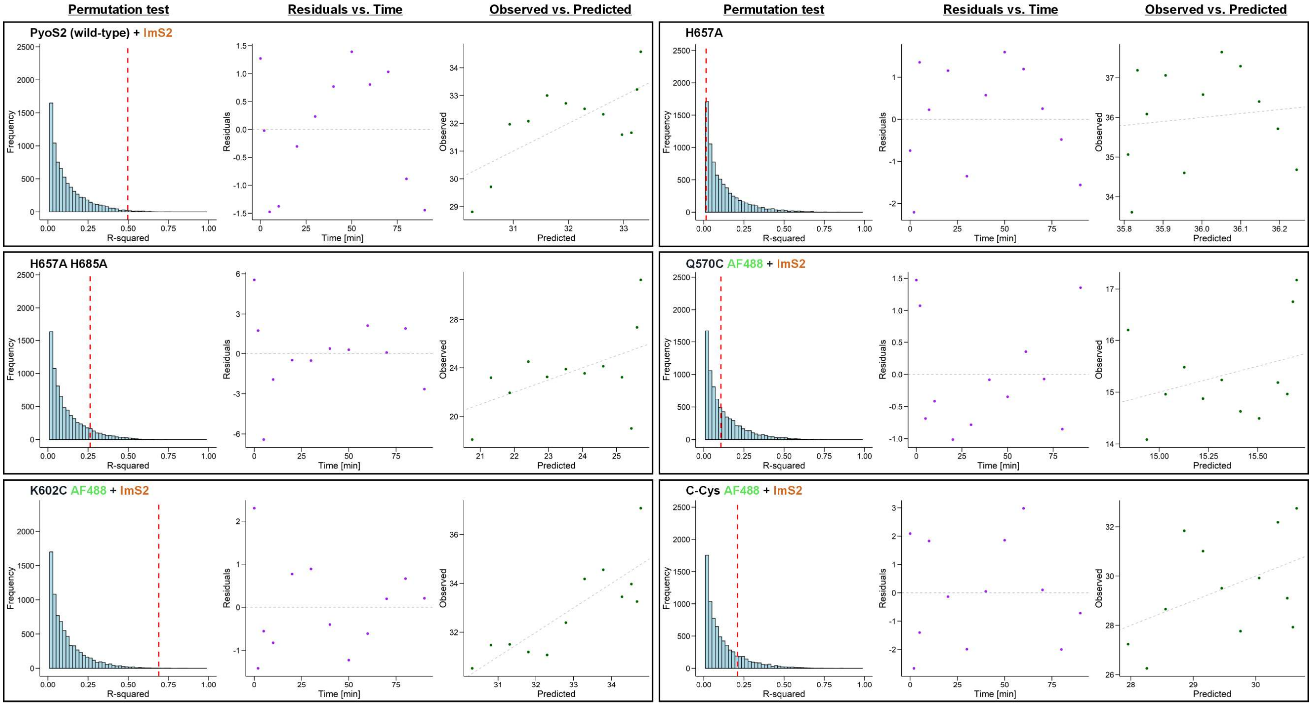


#### Linear regression

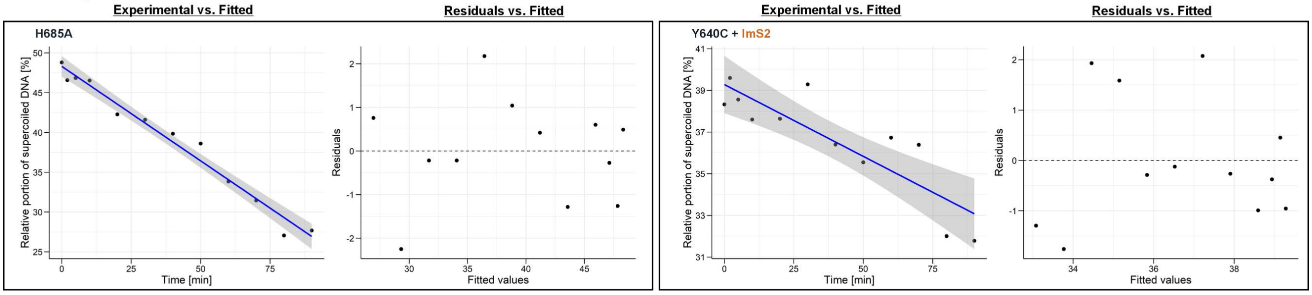


**Supplementary Fig. 5.** Permutation analysis of slow linear DNA accumulation and fitting of supercoiled DNA depletion to a linear regression function. **Top:** Those datasets with a very slow accumulation of linear DNA (data taken from Supplementary Fig. 3) were analysed by permutation analysis to investigate statistical significance from random noise. Shown are three individual plots: i) permutation test (10,000 iterations and $R^2$ highlighted as vertical red line), ii) residuals vs. time, and iii) observed vs. predicted. Fit parameters are summarised in Supplementary Table 5. **Bottom:** The relative supercoiled DNA portion of PyoS2 mutants with **very low** DNase activity (data taken from Supplementary Fig. 3) is fitted to a linear regression curve. Shown are two individual plots: i) experimental data vs. fitted data (including the 95% confidence interval) and ii) residuals vs. fitted. Fit parameters and the estimated half-life of supercoiled DNA are summarised in Supplementary Table 6.

**Supplementary Table 4.** Fit data of **supercoiled** DNA degradation by different variants and conjugates of PyoS2 with apparent exponential decline (very high DNase activity). Curves of relative supercoiled DNA portions from densitometric analysis were fitted to a stretched exponential decay function (equation 2.1) for the following PyoS2 variants. Fits were generated with R.

| | | **Exponential decay – Fit parameters** | | | | | | | | | | | | |
|---|---|---|---|---|---|---|---|---|---|---|---|---|---|---|
| **PyoS2 variant** | **ImS2** | **$y_0$ [%]** | **$y_0$ (lower) [%]** | **$y_0$ (upper) [%]** | **k $[min]^{-1}$** | **k (lower) $[min]^{-1}$** | **k (upper) $[min]^{-1}$** | **b** | **b (lower)** | **b (upper)** | **$R^2$** | **RSE** | **converged** | **$t_{1/2}$ [min]** |
| wild-type | – | 43.14 | 39.90 | 46.36 | 0.26 | 0.22 | 0.30 | 1.46 | 1.04 | NA | 0.99 | 1.43 | TRUE | 3.20 |
| Q570C AF488 | – | 41.61 | 38.65 | 44.59 | 0.29 | 0.25 | 0.34 | 1.11 | 0.81 | 1.59 | 0.99 | 1.32 | TRUE | 2.65 |
| K602C AF488 | – | 42.04 | 39.35 | 44.87 | 0.06 | 0.05 | 0.06 | 1.13 | 0.95 | 1.38 | 0.99 | 1.40 | TRUE | 13.62 |
| Y640C | – | 41.21 | 39.83 | 42.63 | 0.05 | 0.05 | 0.06 | 1.68 | 1.47 | 1.93 | 1.00 | 0.83 | TRUE | 15.26 |
| Y640C AF350 | – | 40.46 | 40.24 | 40.68 | 0.27 | 0.26 | 0.27 | 0.99 | 0.97 | 1.01 | 1.00 | 0.10 | TRUE | 2.80 |
| Y640C AF350 | +ImS2 | 42.25 | 41.18 | 43.32 | 0.11 | 0.10 | 0.11 | 1.14 | 1.05 | 1.24 | 1.00 | 0.50 | TRUE | 7.19 |
| Y640C AF488 | – | 43.36 | 42.83 | 43.89 | 0.19 | 0.18 | 0.19 | 1.13 | 1.09 | 1.18 | 1.00 | 0.24 | TRUE | 4.00 |
| Y640C AF488 | +ImS2 | 38.53 | 36.96 | 40.17 | 0.04 | 0.04 | 0.04 | 1.17 | 1.04 | 1.32 | 1.00 | 0.89 | TRUE | 18.06 |
| Y640C AF594 | – | 40.46 | 39.22 | 41.71 | 0.19 | 0.18 | 0.20 | 0.99 | 0.89 | 1.11 | 1.00 | 0.56 | TRUE | 3.86 |
| Y640C AF594 | +ImS2 | 41.12 | 39.22 | 43.07 | 0.06 | 0.06 | 0.07 | 1.06 | 0.94 | 1.22 | 1.00 | 0.97 | TRUE | 11.54 |
| C-Cys AF488 | – | 38.87 | 36.35 | 41.54 | 0.05 | 0.04 | 0.06 | 1.16 | 0.96 | 1.43 | 0.99 | 1.38 | TRUE | 15.11 |

Abbreviations: $DNA_L$ – linear DNA form; $DNA_S$ – supercoiled DNA form; NA – not available; RSE – residual standard error.

**Supplementary Table 5.** Permutation analysis of slow **linear** DNA accumulation in PyoS2 variants with putative (very low) DNase activity (is the slow increase in linear DNA statistically significantly different from random noise?). Permutations were performed in R (10,000 permutations) and $R^2$ calculated according to equation 5. A p value < 0.05 refers to the 95% confidence interval.

| | | **Permutation analysis – Fit parameters** | | |
|---|---|---|---|---|
| **PyoS2 variant** | **ImS2** | **$R^2$** | **p value** | **interpretation** |
| wild-type | +ImS2 | 0.50 | $7.70 \times 10^{-3}$ | Moderate congruency between data and linear fit; statistically significant (**no** DNase activity) |
| H657A | – | 0.015 | 0.70 | Very low congruency between data and linear fit (**no** DNase activity) |
| H657A H685A | – | 0.26 | $8.55 \times 10^{-2}$ | Low congruency between data and linear fit; statistically significant (**no** DNase activity) |
| Q570C AF488 | +ImS2 | 0.11 | 0.30 | Low congruency between data and linear fit; statistically not significant (**no** DNase activity) |
| K602C AF488 | +ImS2 | 0.69 | $1.40 \times 10^{-3}$ | Moderate congruency between data and linear fit; statistically significant (**no** DNase activity) |
| C-Cys AF488 | +ImS2 | 0.21 | 0.13 | Low congruency between data and linear fit; statistically not significant (**no** DNase activity) |

**Supplementary Table 6.** Fit data of **supercoiled** DNA degradation by different variants and conjugates of PyoS2 with apparent linear decline (very low DNase activity). Curves of relative supercoiled DNA portions from densitometric analysis were fitted to a linear fit function (equation 3.1) for the following PyoS2 variants. Fits were generated with R. A p value < 0.05 refers to the 95% confidence interval.

| | | **Linear regression – Fit parameters** | | | | | | | | |
|---|---|---|---|---|---|---|---|---|---|---|
| **PyoS2 variant** | **ImS2** | **intercept $y_0$ [%]** | **slope b [%/min]** | **$R^2$** | **adjusted $R^2$** | **Std_error** | **t value** | **p value** | **F statistic** | **$t_{1/2}$ [min]** |
| H685A | – | 48.32 | –0.24 | 0.98 | 0.97 | 0.01 | –20.04 | $2.11 \times 10^{-9}$ | 401.48 | 100.66 |
| Y640C | +ImS2 | 39.28 | –0.07 | 0.75 | 0.72 | 0.01 | –5.46 | $2.79 \times 10^{-4}$ | 29.76 | 291.94 |

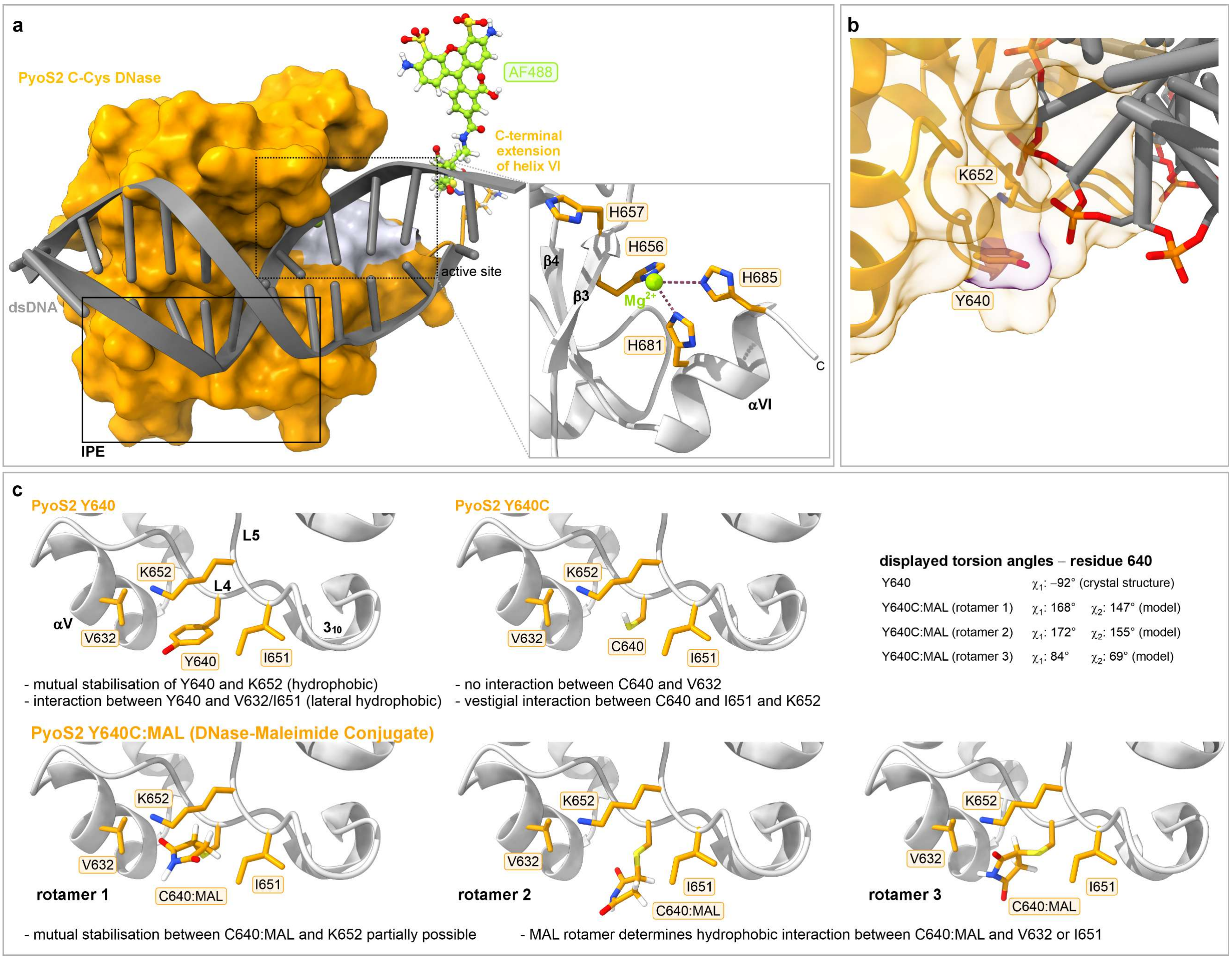


**Supplementary Fig. 6.** Snapshots of the PyoS2 DNase explaining observed effects of AF-labelling on DNase activity. **a**. Manual build of the AF488 fluorophore protruding from helix VI of the PyoS2 crystal structure (4QKO). The C-terminal residues 'GGK' omitted in the crystal structure and the additionally introduced C-terminal cysteine were added manually. **b**. Y640 is not directly interacting with the dsDNA substrate but is in close proximity to it, explaining why maleimide labelling with bulky fluorophores (e.g. AF594) negatively affects DNA binding and thereby reduces DNase activity in comparison to the smaller PyoS2 Y640C AF350 conjugate. **c**. Excerpt of PyoS2's IPE with the original Y640 residue, the unlabelled Y640C substitution, and maleimide-labelled Y640C:MAL (3 rotamers shown, no fluorophore attached), highlighting the mutual stabilisation of Y640 and K652 in PyoS2 and the partial restoration of intraDNase contacts upon maleimide labelling.

**Supplementary Table 7.** List of 113 MDR clinical isolates of *P. aeruginosa* tested by multiplex PCR for *fpvA* subtypes and *pyoS2* genotype. Positive results for *fpvAI*, *fpvAII*, *fpvAIII*, unknown subtypes, and *pyoS2_imS2*⁺ genotypes are indicated in the isolate No. column.

| Isolate No. | Resistance Gene | Date of Isolation | Origin of Specimen | Isolate No. | Resistance Gene | Date of Isolation | Origin of Specimen |
|---|---|---|---|---|---|---|---|
| M0 | GIM-1 | 2002-03-01 | tracheal secretion | M144 | VIM-2 | 2015-05-19 | nasopharyngeal scr. |
| M1 | GIM-1 | 2007-12-09 | tracheal secretion | M145 | GIM-1 | 2015-06-03 | stool |
| M2 | GIM-1 | 2008-01-18 | stool | M146 | GIM-1 | 2015-07-06 | stool |
| M3 | GIM-1 | 2008-04-15 | tracheal secretion | M147 | VIM-2 | 2015-07-08 | tracheal secretion |
| M4 | GIM-1 | 2008-04-13 | urine | M149 | GIM-1 | 2015-08-11 | urine |
| M5 | GIM-1 | 2008-06-04 | tracheal secretion | M150 | GIM-1 | 2015-08-18 | tracheal secretion |
| M7 | GIM-1 | 2008-09-14 | tracheal secretion | M151 | GIM-1 | 2015-08-17 | nasopharyngeal scr. |
| M8 | GIM-1 | 2008-09-22 | tracheal secretion | M152 | GIM-1 | 2015-09-01 | nasopharyngeal scr. |
| M10 | GIM-1 | 2008-10-10 | stool | M159 | GIM-1 | 2015-10-28 | urine |
| M16 | GIM-1 | 2010-02-07 | tracheal secretion | M165 | VIM-2 | 2015-12-29 | tracheal secretion |
| M19 | GIM-1 | 2007-10-10 | wound swab | M166 | GIM-1 | 2016-01-04 | stool |
| M25 | GIM-1 | 2010-02-23 | urine | M171 | GES | 2016-01-28 | catheter stent |
| M28 | GIM-1 | 2010-05-10 | urine | M174 | GES | 2016-03-23 | wound swab |
| M29 | GIM-1 | 2010-05-11 | tracheal secretion | M182 | GIM-1 | 2016-07-07 | drainage secretion |
| M45 | GIM-1 | 2010-11-07 | tracheal secretion | M189 | GIM-1 | 2016-10-07 | abdominal swab |
| M46 | GIM-1 | 2010-12-07 | tracheal secretion | M190 | none | 2016-10-25 | tracheal secretion |
| M47 | GIM-1 | 2010-12-09 | urine | M191 | GIM-1 | 2016-10-25 | nasopharyngeal scr. |
| M48 | GIM-1 | 2011-03-30 | wound swab | M195 | VIM-2 | 2017-01-10 | brain tissue |
| M49 | GIM-1 | 2011-05-31 | tracheal secretion | M199 | GIM-1 | 2017-02-22 | urine |
| M50 | GIM-1 | 2011-06-16 | urine | M206 | VIM-2 | 2017-04-09 | scr. pharynx |
| M51 | GIM-1 | 2011-06-19 | tracheal secretion | M207 | VIM-2 | 2017-04-09 | tracheal secretion |
| M52 | GIM-1 | 2011-06-21 | tracheal secretion | M208 | VIM-2 | 2017-08-18 | tracheal secretion |
| M53 | GIM-1 | 2011-06-21 | tracheal secretion | M209 | GIM-1 | 2017-08-18 | tissue |
| M55 | GIM-1 | 2011-07-03 | bronchoalveolar lavage | M210 | GIM-1 | 2017-09-25 | tracheal secretion |
| M57 | GIM-1 | 2011-04-05 | tracheal secretion | M214 | GIM-1 | 2017-10-24 | catheter urine |
| M58 | GIM-1 | 2011-09-26 | wound swab | M219 | VIM-2 | 2017-12-04 | swab |
| M59 | VIM-2 | 2011-10-10 | tracheal secretion | M224 | GIM-1 | 2018-05-08 | sputum |
| M63 | GIM-1 | 2011-12-25 | tracheal secretion | M228 | VIM-2 | 2018-09-12 | scr. axilla/inguinal region |
| M64 | GIM-1 | 2012-01-23 | wound swab | M230 | VIM-2 | 2018-09-19 | swab |
| M67 | GIM-1 | 2012-03-16 | wound swab | M238 | GIM-1 | 2018-10-31 | bronchial secretion |
| M68 | GIM-1 | 2012-03-19 | urine | M249 | GIM-1 | 2019-01-23 | blood culture |
| M69 | GIM-1 | 2012-04-02 | wound swab | M259 | VIM-2 | 2019-03-13 | central venous catheter |
| M70 | GIM-1 | 2012-05-07 | tracheal secretion | M263 | VIM-2 | 2019-06-11 | blood culture |
| M71 | GIM-1 | 2012-05-21 | pleural aspirate | M273 | GIM-1 | 2019-09-12 | peritoneal swab |
| M72 | GIM-1 | 2012-06-04 | central venous catheter | M283 | GIM-1 | 2019-10-29 | rectal swab |
| M74 | VIM-1 | 2012-08-22 | swab | M287 | VIM-2 | 2019-11-28 | tracheal secretion |
| M78 | VIM-2 | 2013-02-01 | urine | M292 | VIM-2 | 2020-01-09 | urinary catheter |
| M81 | GIM-1 | 2013-03-18 | urine | M294 | GIM-1 | 2020-01-16 | swab anal |
| M88 | GIM-1 | 2013-05-19 | tracheal secretion | M303 | none | 2020-03-05 | axilla/inguinal swab |
| M91 | VIM-2 | 2013-07-12 | urine | M304 | GIM-1 | 2020-03-05 | axilla/inguinal swab |
| M98 | VIM-2 | 2013-09-30 | tracheal swab | M305 | GES | 2020-03-26 | urine |
| M100 | VIM-2 | 2013-10-25 | urine | M307 | GIM-1 | 2020-04-09 | axilla/inguinal swab |
| M103 | VIM-2 | 2013-11-15 | central venous catheter | M313 | VIM-2 | 2020-06-24 | axilla/inguinal swab |
| M105 | IMP-15 | 2013-12-06 | bronchial secretion | M314 | GIM-1 | 2020-06-24 | tracheal secretion |
| M106 | VIM-2 | 2014-01-10 | urethra | M315 | GIM-1 | 2020-07-13 | tracheal secretion |
| M108 | VIM-2 | 2014-02-27 | stool | M320 | VIM-2 | 2020-08-06 | axilla/inguinal swab |
| M119 | GIM-1 | 2014-07-08 | scr. anal | M322 | VIM-2 | 2020-08-12 | blood culture |
| M120 | GIM-1 | 2014-07-09 | tracheal secretion | M323 | GIM-1 | 2020-08-26 | rectal swab |
| M123 | GIM-1 | 2014-07-31 | sputum | M324 | GIM-1 | 2020-09-09 | intra-operative swab |
| M126 | VIM-2 | 2014-08-13 | urine | M325 | VIM-2 | 2020-09-09 | urine |
| M127 | VIM-2 | 2014-09-03 | scr. anal | M326 | GIM-1 | 2020-09-09 | rectal swab |
| M128 | VIM-2 | 2014-09-16 | stool | M328 | GIM-1 | 2020-09-30 | stool |
| M132 | VIM-2 | 2014-10-23 | tracheal secretion | M331 | GIM-1 | 2020-11-05 | unknown |
| M134 | VIM-2 | 2015-11-13 | drainage aspiration | M334 | VIM-2 | 2020-11-25 | nasopharyngeal swab |
| M138 | VIM-2 | 2014-12-22 | scr. anal | M335 | VIM-2 | 2020-12-02 | axilla/inguinal swab |
| M139 | VIM-2 | 2015-02-25 | tracheal secretion | | | | |
| M142 | GIM-1 | 2015-04-21 | nasopharyngeal scr. | | | | |
| M143 | GIM-1 | 2015-04-22 | sputum | | | | |

Abbreviations: scr – screening; GIM-1 – German imipenemase 1; VIM-1 – Verona integron-encoded metallo-β-lactamase 1; VIM-2 – Verona integron-encoded metallo-β-lactamase 2; GES – Guiana extended-spectrum β-lactamase; IMP-15 – Imipenemase 15.

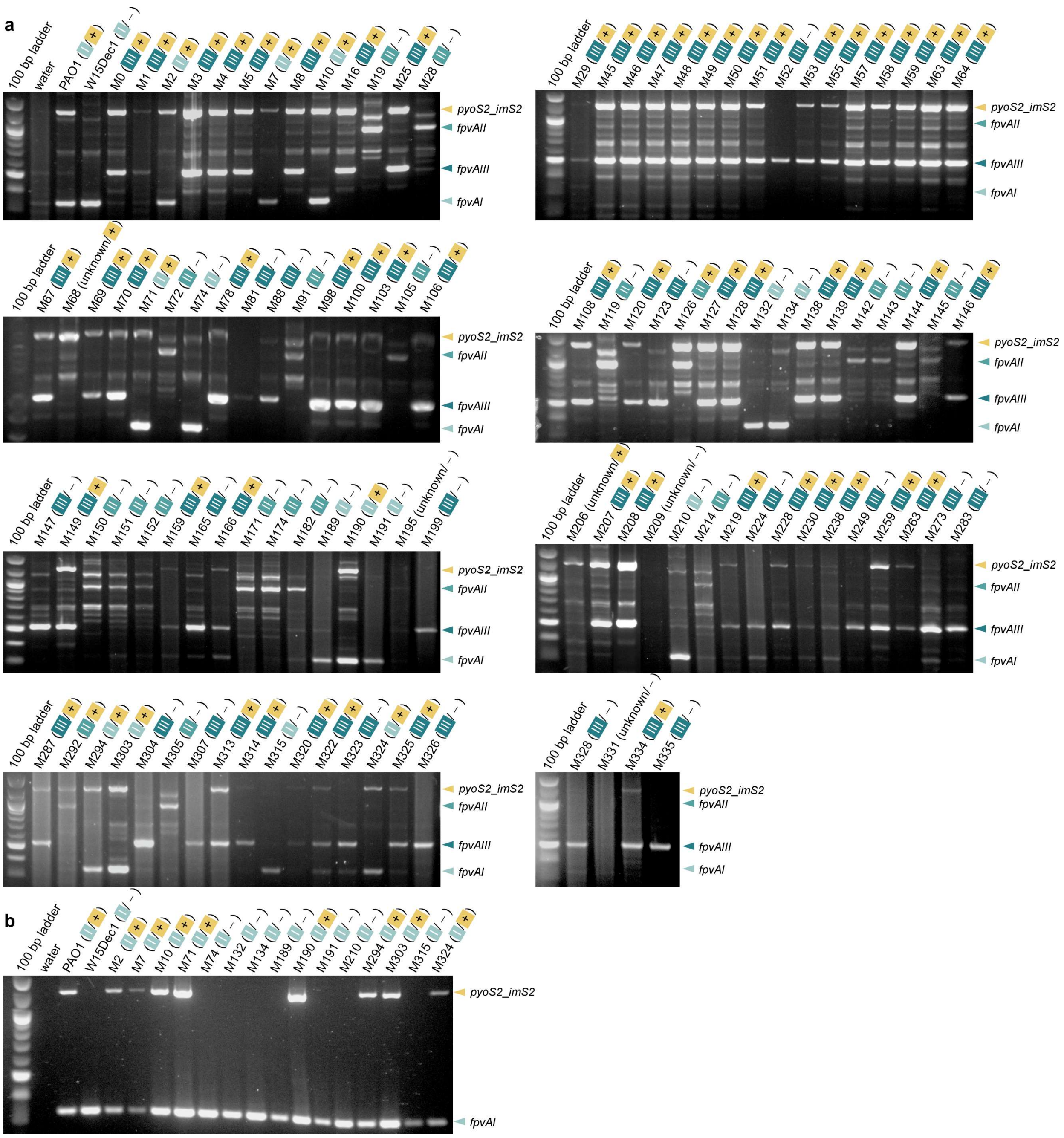


**Supplementary Fig. 7**. Multiplex PCR typing of CRPsA isolates for determining the *fpvA* siderovar and identifying PyoS2 producers. **a**. Agarose gels from multiplex PCR of CRPsA strains from the University Hospital Düsseldorf for detecting amplicons for *fpvAI* (326 bp), *fpvAII* (897 bp), *fpvAIII* (506 bp), and *pyoS2_imS2* (1223 bp) simultaneously. **b**. Agarose gels from duplex PCR of apparently *fpvAI*-positive strains with the primer pairs FpvAI-F/R and PyoS2_ImS2-F/R (CRPsA core strain collection). The Quick-Load 100 bp DNA ladder (New England Biolabs) is used as a size reference. The identified *fpvA* siderovar for every individual strain is indicated by the corresponding number ( I , II , or III ), followed by the *pyoS2_imS2* genotype [producer ( + ) or non-producer ( – )].

**Supplementary Table 8.** Antibiotic susceptibility of *P. aeruginosa* control strains (PAO1 and W15Dec1) and the core list of MDR clinical CRPsA isolates with *fpvAI* genotype. The minimal inhibitory concentrations (MIC) are given in µg/mL. Strains were either classified according to EUCAST as sensitive (S), resistant (R), resistant with technical uncertainty (*R), insufficient evidence (IE), or sensitive with increased exposure (I).

| **Strain/ Isolate No.** | **MBL resistance gene** | **Antibiotic resistance – MIC [µg/mL]** | | | | | | | | | | |
|---|---|---|---|---|---|---|---|---|---|---|---|---|
| | | **PIP** | **PPT** | **CAZ** | **CPM** | **AZT** | **IPM** | **MEM** | **AMI** | **GEN** | **TOB** | **CIP** |
| **PAO1** | none | <4 (I) | 8 (I) | 2 (I) | 2 (I) | 4 (I) | 2 (I) | 1 (S) | 4 (S) | <1 (IE) | <1 (S) | 0.5 (I) |
| **W15Dec1** | none | 8 (*R) | 16 (*R) | 32 (R) | 16 (R) | 4 (R*) | 2 (I) | <0.25 (S) | 4 (S) | 2 (IE) | >1 (S) | 0.25 (I) |
| **M2** | GIM-1 | >128 (R) | >128 (R) | >64 (R) | >32 (R) | 8 (I) | >16 (R) | >16 (R) | 4 (S) | >16 (IE) | >16 (R) | >4 (R) |
| **M7** | GIM-1 | >128 (R) | >128 (R) | >64 (R) | >32 (R) | 16 (*R) | >16 (R) | >16 (R) | 8 (S) | >16 (IE) | >16 (R) | >4 (R) |
| **M10** | GIM-1 | >128 (R) | >128 (R) | >64 (R) | >32 (R) | 32 (R) | >16 (R) | >16 (R) | 8 (S) | >16 (IE) | >16 (R) | >4 (R) |
| **M71** | GIM-1 | >128 (R) | >128 (R) | >64 (R) | >32 (R) | 8 (I) | >16 (R) | >16 (R) | 4 (S) | >16 (IE) | >16 (R) | >4 (R) |
| **M74** | VIM-1 | >128 (R) | >128 (R) | >64 (R) | >32 (R) | 4 (I) | >16 (R) | >16 (R) | 8 (S) | >16 (IE) | >16 (R) | 0.25 (I) |
| **M132** | VIM-2 | >128 (R) | >128 (R) | >64 (R) | >32 (R) | 8 (I) | >16 (R) | >16 (R) | 4 (S) | >16 (IE) | >16 (R) | >4 (R) |
| **M134** | VIM-2 | >128 (R) | >128 (R) | >64 (R) | >32 (R) | 16 (*R) | >16 (R) | 8 (I) | 2 (S) | >16 (IE) | >16 (R) | 0.5 (I) |
| **M189** | GIM-1 | >128 (R) | >128 (R) | >64 (R) | 16 (R) | 4 (*R) | >16 (R) | >16 (R) | 2 (S) | 8 (IE) | >16 (R) | >4 (R) |
| **M190** | none | >128 (R) | >128 (R) | 32 (R) | 16 (R) | 16 (*R) | >16 (R) | 8 (I) | 8 (S) | 4 (IE) | <1 (S) | >4 (R) |
| **M191** | GIM-1 | >128 (R) | >128 (R) | >64 (R) | 16 (R) | 16 (*R) | >16 (R) | >16 (R) | 2 (S) | >16 (IE) | >16 (R) | >4 (R) |
| **M210** | GIM-1 | >128 (R) | >128 (R) | >64 (R) | >32 (R) | 32 (R) | >16 (R) | >16 (R) | 4 (S) | >16 (IE) | >16 (R) | >4 (R) |
| **M294** | GIM-1 | >128 (R) | >128 (R) | >64 (R) | >32 (R) | >64 (R) | >16 (R) | >16 (R) | 4 (S) | >16 (IE) | >16 (R) | >4 (R) |
| **M303** | none | >128 (R) | >128 (R) | >64 (R) | 16 (R) | 16 (*R) | >16 (R) | >16 (R) | 2 (S) | <1 (IE) | <1 (S) | 0.25 (I) |
| **M315** | GIM-1 | >128 (R) | >128 (R) | >64 (R) | 16 (R) | 16 (*R) | >16 (R) | >16 (R) | 4 (S) | >16 (IE) | >16 (R) | >4 (R) |
| **M324** | GIM-1 | >128 (R) | >128 (R) | >64 (R) | >32 (R) | >64 (R) | >16 (R) | >16 (R) | 4 (S) | >16 (IE) | >16 (R) | >4 (R) |

Abbreviations for antibiotics: AMI – amikacin; AZT – aztreonam; CAZ – ceftazidime; CIP – ciprofloxacin; CPM – cefepime; GEN – gentamicin; IPM – imipenem; MEM – meropenem; PIP – piperacillin; PPT – piperacillin/tazobactam; TOB – tobramycin. Abbreviations for resistance genes: GIM-1 – German imipenemase 1; VIM-1 – Verona integron-encoded metallo-β-lactamase 1; VIM-2 – Verona integron-encoded metallo-β-lactamase 2.

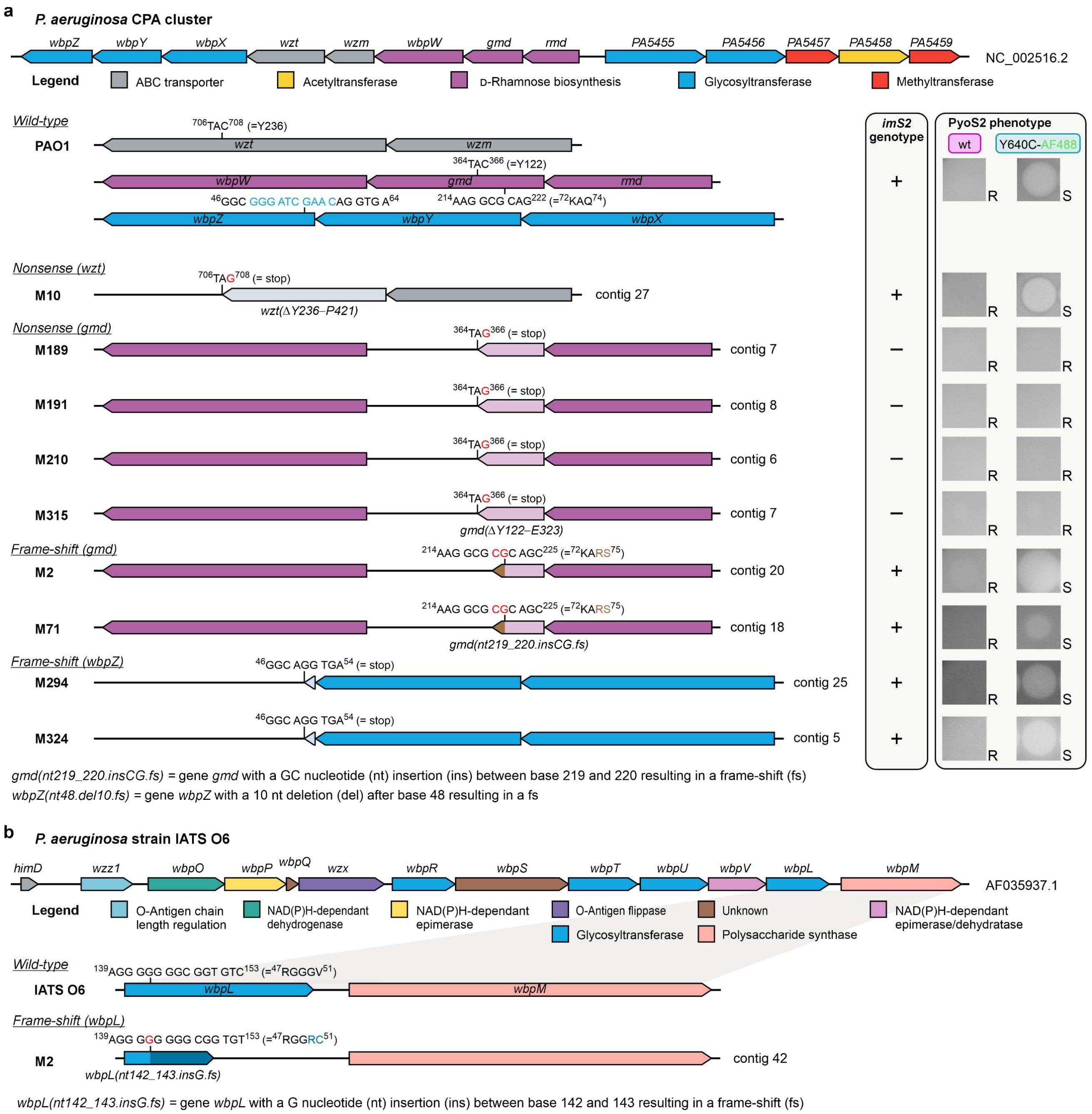


**Supplementary Fig. 8.** Identified LPS mutations in CRPsA isolates sharing the *fpvAI* genotype. In consequence of mutations in the *gmd*, *wzt*, and *wbpZ* locus, the secondary PyoS2 receptor CPA is no longer displayed on the cell surface. The loss of CPA on the cell surface compromises the cytotoxic activity of PyoS2 Y640C variants. A frame-shift mutation in the *wbpL* locus of the O6 serotype results in LPS-free cells, neither displaying CPA nor the O6 OSA on the cell surface. All described mutations lead to pseudogenes.

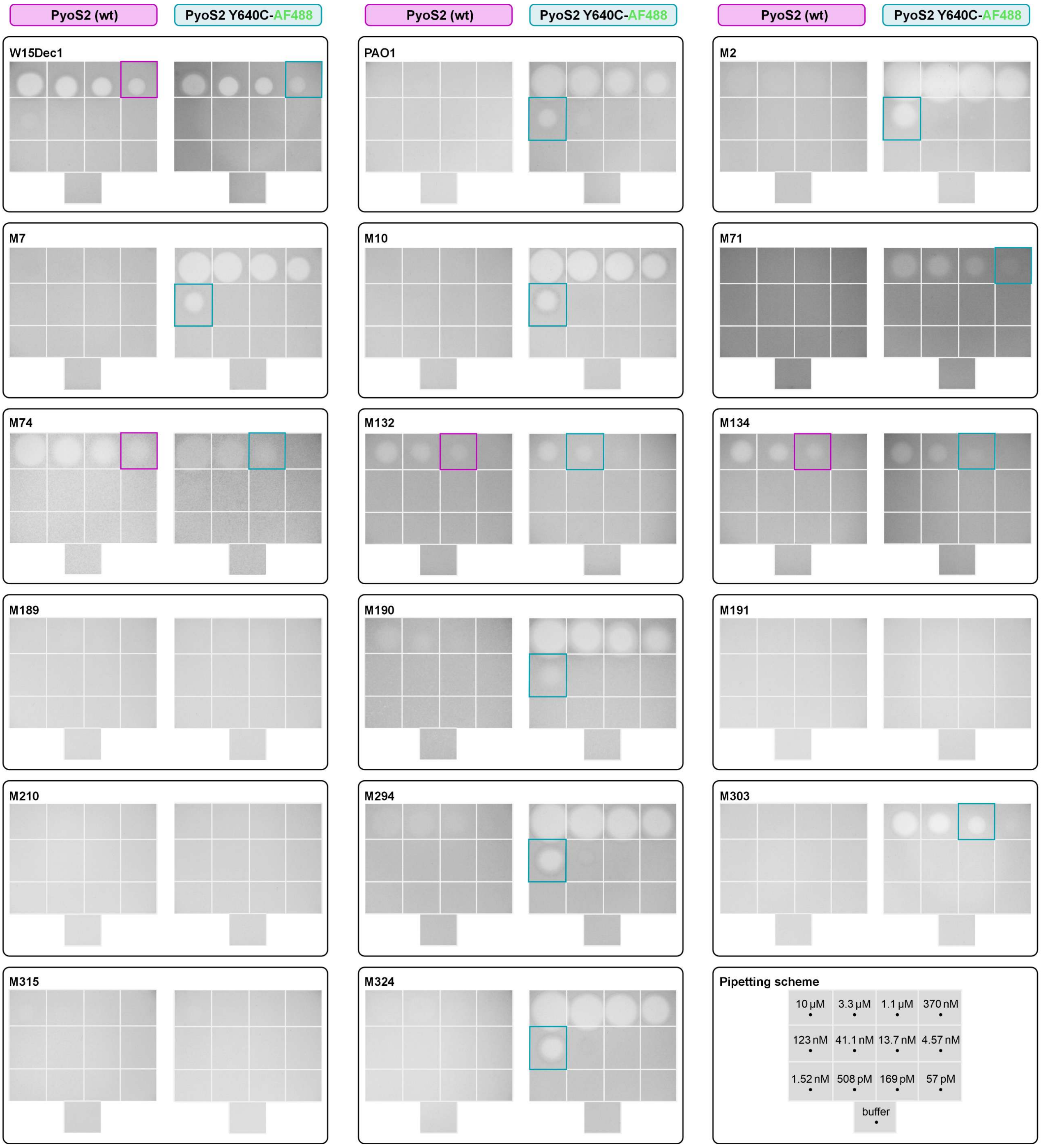


**Supplementary Fig. 9.** Determination of minimal inhibitory concentrations (MICs) in a spot dilution assay. MICs of PyoS2 (wild-type) and the PyoS2 Y640C AF488 conjugate were determined by spot dilution assays on immobilised *P. aeruginosa* strains W15Dec1, PAO1, and the CRPsA isolates M2–M324. Toxin dilutions (5 µL) were used in a range between 10 µM and 57 pM (see pipetting scheme). Pink rectangles highlight the determined MIC for the wild-type toxin, turquoise rectangles those for the PyoS2 Y640C AF488 conjugate. Where no MIC is highlighted, strains showed resistance to concentrations >10 µM. Clear spots indicate antibacterial activity.

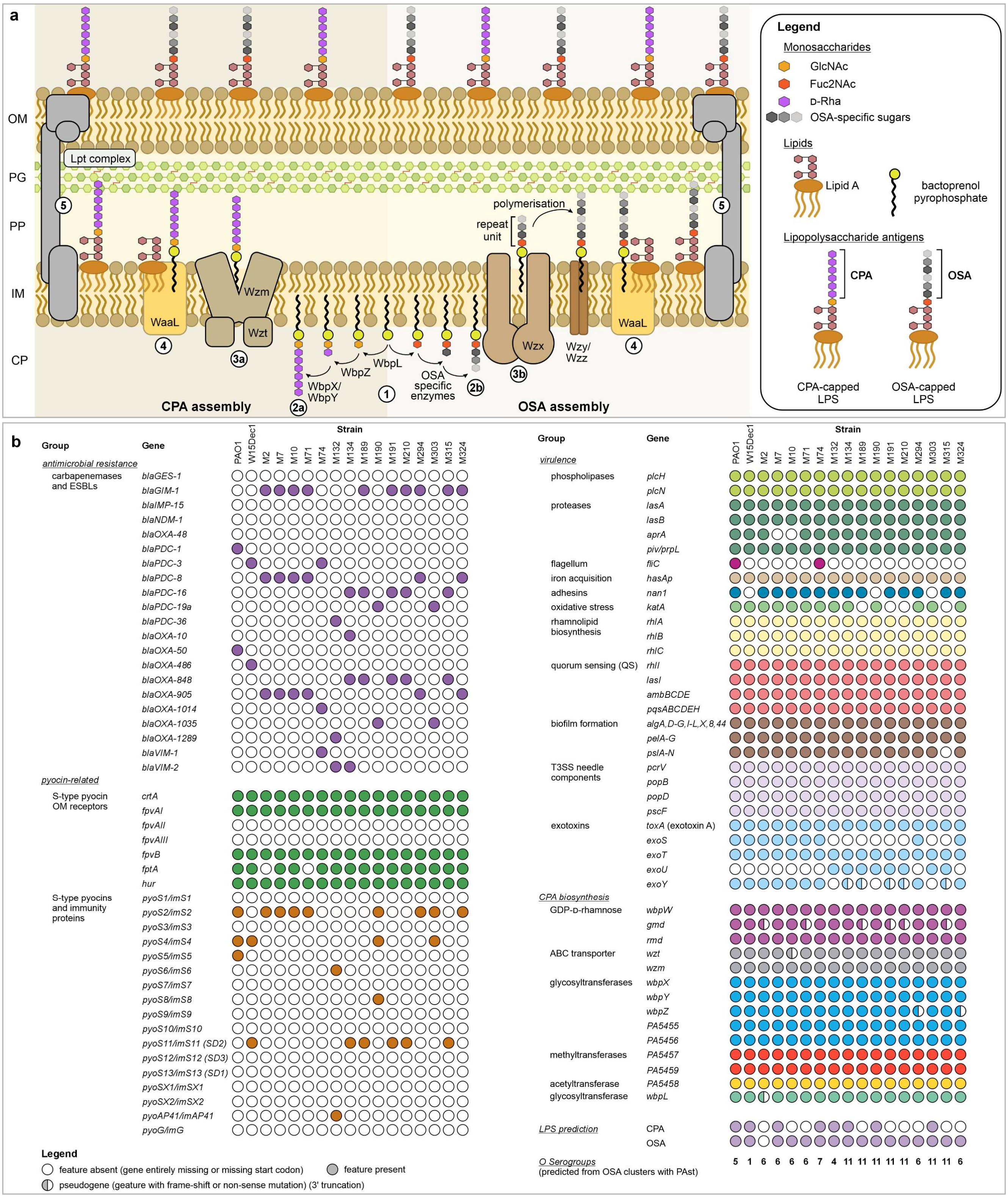


**Supplementary Fig. 10. a**. Assembly pathways for the CPA and OSA antigens of *P. aeruginosa* PAO1 (O5 serotype). The CPA is assembled in an ABC transporter-dependent pathway, while the OSA ligation involves the flippase/translocase Wzx. The synthesis of both antigens is initiated by the transfer of GlcNAc (CPA) or Fuc2NAc (OSA) onto membrane-anchored bactoprenol pyrophosphate, catalysed by the glycosyltransferase WbpL (**1**). While the CPA antigen is fully assembled in the cytoplasm (**2a**) and brought to the periplasmic leaflet of the IM via the ABC transporter Wzm/Wzt (**3a**), the OSA is synthesised in the cytoplasm in small repeating units (**2b**), which are then polymerised in the periplasm through the action of the polysaccharide polymerases Wzy and Wzz (**3b**). The LPS ligase WaaL transfers the CPA and OSA antigen onto lipid A (**4**) and the fully assembled and capped LPS molecules are brought to the cell surface through the membrane-spanning Lpt complex (**5**). Abbreviations: CP – cytoplasm; IM – inner membrane; OM – outer membrane; PG – peptidoglycan; PP – periplasm. **b**. Overview of antibiotic resistance genes, S-type pyocins, common virulence factors, and CPA biosynthesis genes among the *fpvAI*-positive core strain collection of CRPsA strains.

#### Materials and methods

##### Cultivation media and bacterial cultivation

Bacterial strains were cultivated using standard microbiological techniques. For cultivation in liquid culture, bacterial strains were either grown in lysogeny broth (LB) (10 g/L tryptone/peptone, 5 g/L yeast extract, 10 g/L NaCl) or terrific broth (TB) [50.8 g/L TB mix (Carl Roth, Germany), 4 mL/L glycerol] media, both supplemented with 100 µg/mL ampicillin (all pET21 vectors) and 25 µg/mL chloramphenicol [only *E. coli* Rosetta 2(DE3) pLysS] if required. Furthermore, *E. coli* strains were plated on LB agar (1.4% agar-agar) for microbial selection. For cloning and protein production, chemically competent *E. coli* host strains were transformed with plasmid DNA using the heat-shock procedure. For CRPsA sampling, *P. aeruginosa* strains were plated on selective and non-selective media, e.g. Columbia agar with 5% sheep blood (Biomérieux, Germany), MacConkey agar (Biomérieux, Germany), and ESBL agar (with ceftazidime or cefotaxime). All bacterial strains used in this work are summarised in Supplementary Table 9.

**Supplementary Table 9.** Bacterial strains used in this work.

| Strain | Genotype or relevant characteristic | Reference |
|---|---|---|
| | ***Pseudomonas aeruginosa*** | |
| PAO1 | Clinical isolate, burn wound, PyoS2-resistant, ATCC 15692 | [19] |
| W15Dec1 | Environmental isolate from a Belgian river, PyoS2-sensitive | [20] |
| M strains | MDR clinical isolates from hospitalised patients (for details see Supplementary Table 7) | this work |
| | ***Escherichia coli*** | |
| DH5α | F⁻ *endA1 glnV44 thi-1 recA1 relA1 gyrA96 deoR nupG purB20* φ80*dlacZ*ΔM15 *Δ(lacZYA-argF)*U169, hsdR17($r_K^-$ $m_K^+$), $\lambda^-$ | [21] |
| BL21(DE3) | *E. coli* strain B, F⁻ *ompT gal dcm lon* hsdSB($r_B^-$ $m_B^-$) *λ*(DE3) [*lacI lacUV5-T7p07 ind1 sam7 nin5*]) [*malB*⁺]$_{K-12}$($\lambda^S$) | [22] |
| C43(DE3) | *E. coli* strain BL21, F⁻ *ompT gal dcm* hsdSB($r_B^-$ $m_B^-$) *λ*(DE3), uncharacterised mutations confer resistance to toxic proteins | [23] |
| JM109(DE3) | *endA1 recA1 gyrA96 thi hsdR17(rk⁻ mk⁺), relA1, supE44, λ⁻, Δ(lac-proAB),* [*F', traD36, proAB, lacI$^q$ZΔM15*], *λ*(DE3) | [24] |
| Rosetta 2(DE3) pLysS | F⁻ *ompT hsdSB*($r_B^-$ $m_B^-$) *gal dcm* pRARE2 *(Cam$^R$)* pLysS (*Cam*$^R$) | Novagen |

##### DNA oligonucleotides

DNA oligonucleotides were ordered from Sigma Aldrich (Merck) in quantities of 0.025 µmol (desalted). Upon arrival, all oligos were dissolved in deionised water (100 µM) and stored at –20 °C. They were used as primers for DNA sequencing according to Sanger, for PCR amplification of genes in the process of molecular cloning, for PCR-based multiplex detection of siderophore subtypes in *P. aeruginosa*, or as molecular probes for MBL genes in MDR clinical isolates of *P. aeruginosa*. All DNA oligonucleotides used throughout this work are summarised in Supplementary Table 10 and 11.

**Supplementary Table 10.** DNA oligonucleotides for cloning of PyoS2 derivatives, Sanger sequencing, and multiplex PCR characterisation in MDR isolates of *P. aeruginosa* regarding their *fpvA* subtypes (this work) and MBL genes (at UKD Düsseldorf).

| Primer | Sequence (5'-3') | Purpose |
|---|---|---|
| | **Cloning primers** | |
| TWE01 | TATGGCTAGCATGGCTGTCAATGATTACG | *pyoS2_imS2-His$_6$* restrict-ion cloning into pET21a(+) |
| TWE02 | TAATACTCGAGACCGGCCTTAAAGCCAGGAAGC | |
| TWE09 | GGAAATCTTGGACTTCATATCAACACTTCCCTCCCTTGTGGATTTC | addition of a C-terminal cysteine residue to PyoS2 |
| TWE10 | GAAATCCACAAGGGAGGGAAGTGTTGATATGAAGTCCAAGATTTCC | |
| TWE17 | CGTAATCATTGACAGCCATATGTATATCTCCTTCTTAAAG | deletion of 3 N-term. AAs (MAS) from *Nhe*I cloning |
| TWE18 | CTTTAAGAAGGAGATATACATATGGCTGTCAATGATTACG | |
| TWE13 | GAATAAAGATCGAAATCCACGCAAAGGTTCGAATAGCAGATG | H657A substitution |
| TWE14 | CATCTGCTATTCGAACCTTTGCGTGGATTTCGATCTTTATTC | |
| TWE15 | CAAAACGTCATATAGAAATCGCAAAGGGAGGGAAGTGATATG | H685A substitution |
| TWE16 | CATATCACTTCCCTCCCTTTGCGATTTCTATATGACGTTTTG | |
| TWE19 | GAAATCTTGGACTTCATATCAACACTTCCCTCCCTTTGCGATTTC | addition of C-term. cysteine to PyoS2 H657A H685A |
| TWE20 | GAAATCGCAAAGGGAGGGAAGTGTTGATATGAAGTCCAAGATTTC | |
| TWE23 | CCGAAAGTCCCGCCAGTTACAGAATGTCTTACCACGTAG | K602C substitution |
| TWE24 | CTACGTGGTAAGACATTCTGTAACTGGCGGGACTTTCGG | |

| | | |
|---|---|---|
| TWE25 | GAGCCAATTACCGCTGACGGGACATCCCTTGCCAGTCGCAGCACCAG | Q570C substitution |
| TWE26 | CTGGTGCTGCGACTGGCAAGGGATGTCCCGTCAGCGGTAATTGGCTC | |
| TWE29 | CCTGTTCTGACTCTCTGACACAAGGAGCCCCTCCATCTCTCATTACAGC | Y640C substitution |
| TWE30 | GCTGTAATGAGAGATGGAGGGGCTCCTTGTGTCAGAGAGTCAGAACAGG | |
| **Sequencing primers** | | |
| T7 | TAATACGACTCACTATAGGG | for T7-based pET vectors |
| T7term | CTAGTTATTGCTCAGCGGT | for T7-based pET vectors |
| Seq_TWE01 | TAGAGGGAACCGTAACCTCGTAC | for *pyoS2_imS2* inserts |
| Seq_TWE03 | CCGGTAGTGTAATCACCCCAGGATAG | for *pyoS2_imS2* inserts |
| Seq_TWE04 | CGATCTATGTGATGTTCAGGGATC | for *pyoS2_imS2* inserts |

Abbreviation: AA – amino acid.

**Supplementary Table 11**. Multiplex primers for the detection of *fpvA* subtypes, *pyoS2_imS2*, and carbapenemase genes.

| | | |
|---|---|---|
| ***fpvA* and *pyoS2_imS2* multiplex primers for the detection in *P. aeruginosa* genomes** | | |
| FpvAI-F | CGAAGGCCAGAACTACGAGA | *fpvAI* detection |
| FpvAI-R | TGTAGCTGGTGTAGAGGCTCAA | (326 bp fragment) |
| FpvAII-F | TACCTCGACGGCCTGCACAT | *fpvAII* detection |
| FpvAII-R | GAAGGTGAATGGCTTGCCGTA | (897 bp fragment) |
| FpvAIII-F | ACTGGGACAAGATCCAAGAGAC | *fpvAIII* detection |
| FpvAIII-R | CTGGTAGGACGAAATGCGAG | (506 bp fragment) |
| PyoS2_ImS2-F | CAAGCGATCTCCGATGCGATTG | *pyoS2_imS2* detection |
| PyoS2_ImS2-R | CTCCAGCTGGGCTATCTTCTC | (1223 bp fragment) |
| **Carbapenemase multiplex primers and dual-labelled TaqMan probes** | | |
| NDM1-S | 6-FAM-GCGACTTGGCCTTGCTGTCCTTG-BHQ-1 | NDM-1 probe |
| NDM1-F | CGGCATCACCGAGATTGC | NDM-1 forward primer |
| NDM1-R | CACCGACATCGCTTTTGGT | NDM-1 reverse primer |
| VIM-1-S | 6-FAM-CTTCTATCCTGGTGCTGCGCATTCG-BHQ-1 | VIM-1 probe |
| VIM-1-F | TGCGCTTCGGTCCAGTAGA | VIM-1 forward primer |
| VIM-1-R | TGACGGGACGTATACAACCAGAT | VIM-1 reverse primer |
| VIM-2-S | 6-FAM-CTTCTATCCTGGTGCTGCGCATTCG-BHQ-1 | VIM-2 probe |
| VIM-2-F | GCGCTTCGGTCCAGTAGAAC | VIM-2 forward primer |
| VIM-2-R | CTCGCAGACGGGACGTACA | VIM-2 reverse primer |
| KPC-S | 6-FAM-CAGTCGGAGACAAAACCGGAACCTGC-BHQ-1 | KPC probe |
| KPC-F | GCAGCGGCAGCAGTTTGTTGATT | KPC forward primer |
| KPC-R | GTAGACGGCCAACACAATAGGTGC | KPC reverse primer |
| OXA-48-S | Hex-CTGGCTGCGCTCCGATACGTGTAACTTATTG-BHQ-1 | OXA-48 probe |
| OXA-48-F | TTCGGCCACGGAGCAAATCAG | OXA-48 forward primer |
| OXA-48-R | GATGTGGGCATATCCATATTCATCGCA | OXA-48 reverse primer |
| GES-S | TexasRed-CGACCTCAGAGATACAACTACGCCTATTGC-BHQ-2 | GES probe |
| GES-F | CGGTTTCTAGCATCGGGACACAT | GES forward primer |
| GES-R | CCGCCATAGAGGACTTTAGCMACA | GES reverse primer |
| GIM-S | Hex-ACACGAAGTTGTTATTATCCTGGGCGACTGAC-BHQ-1 | GIM-1 probe |
| GIM-F | CGACACACCTTGGTCTGAAGAA | GIM-1 forward primer |
| GIM-R | GATGCTAGCCATAACCTGGTATCC | GIM-1 reverse primer |
| IMP-S | Hex-CGATCTATCCCCACGTATGCATCTGAATTAACA-BHQ-1 | IMP probe |
| IMP-F | GGGCGGAATAGAGTGGCTTA | IMP forward primer |
| IMP-R | GGCTTGAACCTTACCGTCTTTTT | IMP reverse primer |

Abbreviations: BHQ-1 – black hole quencher 1; 6-FAM – 6-carboxyfluorescein; Hex – hexachlorofluorescein.

#### Cloning strategy

The reference sequence of the *pyoS2_imS2* tandem was obtained from the reference genome assembly of *P. aeruginosa* PAO1 (NCBI reference sequence: NC_002516.2)[19]. The corresponding loci *pyS2* (NP_249841.1) and *imm2* (NP_249842.1) were amplified by polymerase chain reaction (PCR), using the primer pair TWE01 and TWE02 (for primer information see Supplementary Table 10) to introduce an *Nhe*I restriction site at the 5'-end of the *pyoS2* gene and an *Xho*I site at the 3'-end of the *imS2* gene. The amplicon was then digested with *Nhe*I and *Xho*I and subsequently ligated with *Nhe*I and *Xho*I linearised pET21a(+) plasmid DNA to give the vector pTWE01 (for plasmid information see Supplementary Table 12). By using the pET21a(+)-backbone, a C-terminal $His_6$-tag was attached to the immunity protein. As the *Nhe*I restriction site contains a second ATG start codon, the artificial three in-frame amino acids MAS on the N-terminus of PyoS2 were removed by QuickChange (QC) PCR with primers TWE17 and TWE18, yielding the desired vector pTWE10, harbouring *pyoS2* and *imS2-His*$_6$. In the following, mutations were introduced into the *pyoS2* sequence by QC PCR to either diminish the DNase activity by substituting the essential $Mg^{2+}$-coordinating His-residues H657 and/or H685 or to enable maleimide labelling on introduced Cys-residues on the C-terminus (C-Cys) or in the positions Q570, K602, or Y640. Based on the wild-type *pyoS2* sequence in pTWE10, the DNase-affected mutants *pyoS2-H657A* (pTWE08), *pyoS2-H685A* (pTWE09), and *pyoS2-H657A-H685A* (pTWE04) were generated. Furthermore, pTWE10 served as a template for the cysteine mutants *pyoS2-Q570C* (pTWE16), *pyoS2-K602C* (pTWE14), *pyoS2-Y640C* (pTWE18), and *pyoS2-C-Cys* (pTWE13). The same approach of cloning was also employed to generate DNase-inactive cysteine mutants from plasmid pTWE04, which encodes for *pyoS2-H657A-H685A*, namely *pyoS2-Q570C-H657A-H685A* (pTWE17), *pyoS2-K602C-H657A-H685A* (pTWE15), *pyoS2-Y640C-H657A-H685A* (pTWE19), and *pyoS2-H657A-H685A-C-Cys* (pTWE05). For microbial selection and plasmid isolation, transformations were typically carried out with chemically competent *E. coli* DH5α. However, since all plasmids harbouring the Y640C mutation displayed a cytotoxic phenotype, these variants were cloned and purified from *E. coli* C43(DE3). All generated plasmids and their inserts were verified by Sanger sequencing at Eurofins Genomics Germany GmbH using the sequencing primers T7, T7term, Seq_TWE01, Seq_TWE03, and Seq_TWE04 as specified in Supplementary Table 10.

**Supplementary Table 12.** Plasmids generated and used in this work.

| Plasmid | Genotype | Encoded Proteins | Template | Reference |
|---|---|---|---|---|
| pET21a(+) | *amp*$^R$, pBR322 Ori, T7 promoter, T7term, *lacI*, *lacO* | None | – | Novagen |
| pTWE01 | pET21a(+), *MAS-pyoS2*, *imS2-His*$_6$ | MAS-PyoS2, ImS2-$His_6$ | PAO1 gDNA | this work |
| pTWE04 | pET21a(+), *pyoS2-H657A-H685A*, *imS2-His*$_6$ | PyoS2 H657A H685A, ImS2-$His_6$ | pTWE08 | this work |
| pTWE05 | pET21a(+), *pyoS2-H657A-H685A-C-Cys*, *imS2-His*$_6$ | PyoS2 H657A H685A C-Cys, ImS2-$His_6$ | pTWE04 | this work |
| pTWE08 | pET21a(+), *pyoS2-H657A*, *imS2-His*$_6$ | PyoS2 H657A, ImS2-$His_6$ | pTWE10 | this work |
| pTWE09 | pET21a(+), *pyoS2-H685A*, *imS2-His*$_6$ | PyoS2 H685A, ImS2-$His_6$ | pTWE10 | this work |
| pTWE10 | pET21a(+), *pyoS2*, *imS2-His*$_6$ | PyoS2, ImS2-$His_6$ | pTWE01 | this work |
| pTWE13 | pET21a(+), *pyoS2-C-Cys*, *imS2-His*$_6$ | PyoS2 C-Cys, ImS2-$His_6$ | pTWE10 | this work |
| pTWE14 | pET21a(+), *pyoS2-K602C*, *imS2-His*$_6$ | PyoS2 K602C, ImS2-$His_6$ | pTWE10 | this work |
| pTWE15 | pET21a(+), *pyoS2-K602C-H657A-H685A*, *imS2-His*$_6$ | PyoS2 K602C H657A H685A, ImS2-$His_6$ | pTWE04 | this work |
| pTWE16 | pET21a(+), *pyoS2-Q570C*, *imS2-His*$_6$ | PyoS2 Q570C, ImS2-$His_6$ | pTWE10 | this work |
| pTWE17 | pET21a(+), *pyoS2-Q570C-H657A-H685A*, *imS2-His*$_6$ | PyoS2 Q570C H657A H685A, ImS2-$His_6$ | pTWE04 | this work |
| pTWE18 | pET21a(+), *pyoS2-Y640C*, *imS2-His*$_6$ | PyoS2 Y640C, ImS2-$His_6$ | pTWE10 | this work |
| pTWE19 | pET21a(+), *pyoS2-Y640C-H657A-H685A*, *imS2-His*$_6$ | PyoS2 Y640C H657A H685A, ImS2-$His_6$ | pTWE04 | this work |

#### Heterologous protein expression

PyoS2 variants were heterologously produced in *E. coli* BL21(DE3) and C43(DE3) [C43 used for all constructs harbouring the Y640C mutation]. Starting with a single colony of the strain carrying the desired plasmid for protein expression (cf. Supplementary Table 12), LB media (100 µg/mL ampicillin) was inoculated and this preculture was incubated overnight at 37 °C (130 RPM). This culture was then grown overnight at 37 °C (130 RPM). 2 L of TB media (100 µg/mL ampicillin) in a 5 L Erlenmeyer flask was inoculated with the overnight culture to an $OD_{600}$ of 0.05 and incubated at 37 °C (130 RPM) until an $OD_{600}$ of 0.6–0.8 was reached. Cells were allowed to chill at RT for 30 min before protein expression was induced by the addition of IPTG to a final concentration of 500 µM. Expression cultures were incubated overnight at 25 °C (130 RPM). Next, cultures were harvested by centrifugation (4,500 RPM, 15 min, 4 °C), the supernatant was discarded, the cell pellets were resuspended in an appropriate amount of binding buffer (20 mM Tris-HCl, 500 mM NaCl, 5 mM imidazole, pH 7.5), and the cell suspensions were stored at –20 °C until needed.

##### Protein purification

After thawing resuspended cells, PMSF protease inhibitor was added to a final concentration of 1 mM (100 mM stock in *i*PrOH). Cells were then disrupted by sonication (3 x 10 min, 40% amplitude, 5 min rest on ice). Cell debris was removed by centrifugation (15,000 x g, 2 x 30 min, 4 °C) and the supernatant was separated from the pellet. Next, the cleared lysate was applied to a Ni-NTA column (Qiagen Ni-NTA Superflow), equilibrated in binding buffer, and afterwards thoroughly washed with binding buffer (min. 40 CV) to remove bulk host proteins and unbound PyoS2/ImS2-$His_6$ complex from the resin (crucial step to avoid ImS2 contamination in PyoS2 samples). All PyoS2 derivatives were purified as previously described for colicins by disrupting the column-bound PyoS2/ImS2-$His_6$ complex by washing with 3 CV of the chaotropic salt-containing complex disruption buffer (20 mM Tris-HCl, 500 mM NaCl, 6 M guanidine hydrochloride, pH 7.5), leaving only the His-tagged ImS2 bound to the NTA resin[25]. After washing the column with 2 CV of binding buffer, the immunity protein was separately eluted with elution buffer (20 mM Tris-HCl, 500 mM NaCl, 250 mM imidazole, pH 7.5). PyoS2-containing and ImS2-containing fractions were merged separately, and EDTA was added to a final concentration of 1 mm. PyoS2-containing fractions were then dialysed overnight at 4 °C against 5 L of dialysis buffer (25 mM Tris-HCl, 150 mM NaCl, pH 8), using 12–14 kDa dialysis tubing (Spectra/Por 4 Dialysis Membrane, Standard RC Tubing, 12–14 kDa MWCO). For ImS2-$His_6$, imidazole from the elution buffer was removed by successive rounds of centrifugal concentration (Sartorius Vivaspin 20, 3 kDa MWCO) and washing with dialysis buffer. All PyoS2 proteins and ImS2-$His_6$ were finally purified by size exclusion chromatography on a Cytiva HiLoad 26/600 Superdex 200 prep grade column with dialysis buffer as isocratic eluent. Monomeric protein species were merged and concentrated to the desired concentration by centrifugal concentration for further usage. Furnished proteins were frozen in liquid $N_2$ and stored at –20 °C until needed.

##### Maleimide labelling of PyoS2 cysteine mutants

Purified cysteine mutants of PyoS2 were thawed on ice and DTT (1 M stock in dialysis buffer) was added to a final concentration of 10 mM to reduce putative disulfide bridges. Incubation was carried out for 60 min on ice. Excess of reducing agent was removed by desalting into labelling buffer (25 mM Tris-HCl, 150 mM NaCl, pH 7.5) on a Cytiva HiLoad desalting column (5 mL). PyoS2-containing fractions were merged and the protein concentration was determined by UV-Vis measurements at 280 nm. The desired fluorophore maleimide (AlphaFluor 350 $C_5$ maleimide, AlphaFluor 488 $C_5$ maleimide, or AlphaFluor 594 $C_5$ maleimide, AAT Bioquest, 10 mM in DMSO) was added to a 3-fold molar excess and labelling was carried out at RT under gentle agitation in the dark for 1.5 h. DTT was added to a final concentration of 5 mM for quenching of excess maleimide and the labelled proteins were subsequently separated from unreacted fluorophore dye by desalting into dialysis buffer (25 mM Tris-HCl, 150 mM NaCl, pH 8.0). After reducing the volume to the desired protein concentration by centrifugal concentration at 4 °C, labelled proteins were stored at –20 °C until needed.

##### Protein quantification and determination of maleimide labelling efficiencies

Purified proteins were quantified by measuring the absorption of aromatic amino acids under UV irradiation at 280 nm according to the Beer-Lambert Law. For this purpose, molar extinction coefficients and molecular weights were predicted and calculated based on the primary amino acid sequence with the Expasy ProtParam tool[26].

After maleimide labelling, PyoS2-AF conjugates were subjected to absorption measurements in dialysis buffer to determine protein concentrations and the labelling efficiency according to the manufacturer (AAT Bioquest). Therefore, the protein absorption was measured at 280 nm and the absorption of the attached AlphaFluor fluorophore dyes was measured at 343 nm (AF350, $\varepsilon_{343}$: 19,000 $M^{-1}$ $cm^{-1}$), 499 nm (AF488, $\varepsilon_{499}$: 71,000 $M^{-1}$ $cm^{-1}$), or 590 nm (AF594, $\varepsilon_{590}$: 90,000 $M^{-1}$ $cm^{-1}$) using a Shimadzu UV-1800 UV spectrophotometer and Hellma Analytics quartz glass cuvette SUPRASIL QS (d = 10 mm light path length). Labelling efficiency was determined as described in the manufacturer's protocol with the following correction factors for the correction of fluorophore absorption at 280 nm: 0.19 (AF350), 0.11 (AF488), and 0.56 (AF594). All fluorescently labelled proteins used for *in vitro* DNase assays and *in vivo* cytotoxicity screening with live cells of *P. aeruginosa* were labelled with greater than 98% efficiency and verified by mass spectroscopy (cf. Supplementary Table 3).

##### Sodium dodecyl sulfate polyacrylamide gel electrophoresis (SDS-PAGE)

Purified proteins were diluted in dialysis buffer (25 mM Tris-HCl, 150 mM NaCl, pH 8.0) to a concentration of 0.33 µg/µL and mixed with 4x SDS sample buffer (200 mM Tris-HCl, pH 6.8, 8% w/v SDS, 0.4% w/v bromophenol blue, 40% w/v glycerol, 400 mM β-mercaptoethanol) to give a final concentration of 0.25 µg/µL. Samples were heated at 98 °C for 10 min and used without centrifugation. 5 µL marker (ThermoScientific PageRuler Unstained Protein Ladder, Ref 26614) and 8 µL protein sample (equals 2 µg protein per well) were applied to a commercial Invitrogen NuPAGE 4−12% Bis-Tris gel (1.0 mm x 15 well, Ref NP0323BOX). Gels were run at 180 V for 46 min in 1x NuPAGE MOPS SDS running buffer (50 mM MOPS, 50 mM Tris base, 0.1% SDS, 1 mM EDTA, pH 7.7, Ref NP0001). After three times washing with water and incubation with colloidal Coomassie o/n, stained gels were incubated in water with a cellulose tissue to destain the background. Gels were then documented by photography.

#### Electrospray ionisation mass spectrometry (ESI-MS)

Purified and labelled PyoS2 and ImS2 proteins were analysed and verified by LC-coupled ESI-MS measurements at the Central Institute of Engineering, Electronics and Analytics (ZEA-3) at the Jülich Research Center. Concentrated samples were diluted 1:10 with water and separated on an Accucore 150-C4 HPLC column (100 mm x 4.6 mm, 2.6 µm particle size), using a 5–95% gradient of acetonitrile. The eluents water and acetonitrile were supplemented with 0.025% heptafluorobutyric acid (HFBA) for ion-pairing and 1% formic acid (FA). Eluent A (water + 0.025% HFBA + 1% FA), eluent B (MeCN + 0.025% HFBA + 1% FA).

#### Multiplex PCR (*fpvA* and *pyoS2_imS2* detection)

A single colony of *P. aeruginosa* MDR clinical isolates and the control strains PAO1 and W15Dec1, grown on COS agar, was suspended in sterile water (50 µL) and heated to 98 °C for 10 min. After centrifugation (20,000 x g, 10 min, 4 °C), the DNA-containing supernatant (DNA template) was transferred to a PCR tube and stored at −20 °C until needed. All necessary ingredients (except 0.4 µL template DNA for multiplex detection or 0.4 µL sterile water for the negative control) were pre-mixed in a master mix and distributed to PCR tubes and furnished with template DNA immediately before the start of the PCR reaction.

**Supplementary Table 13.** Reaction setup for the multiplex PCR detection of *fpvA* genes and the *pyoS2_imS2* tandem.

| | Multiplex [µL] | Control [µL] |
|---|---|---|
| 5x HF buffer | 4 | 4 |
| DMSO | 1.2 | 1.2 |
| 10 mM dNTP mix | 0.4 | 0.4 |
| Primer* (7.5 µM stock in $H_2O$) | 0.4 (for each primer) | 0.4 (for each primer) |
| Sterile water | 10.6 | 11 |
| Phusion polymerase | 0.2 | 0.2 |
| Template DNA | 0.4 | – |
| Total volume | 20 | 20 |

* The multiplex PCR detects four genes at the same time (*fpvAI*, *fpvAII*, *fpvAIII*, and *pyoS2_imS2*). Thus, each PCR reaction contains eight primers: FpvAI-F, FpvAI-R, FpvAII-F, FpvAII-R, FpvAIII-F, FpvAIII-R, PyoS2_ImS2-F, and PyoS2_ImS2-R.

**Supplementary Table 14.** PCR protocol for the amplification of *fpvA* genes and the *pyoS2_imS2* tandem from genomic DNA of *P. aeruginosa* strains.

| | Temperature [°C] | Time | |
|---|---|---|---|
| Initial denaturation | 98 | 2 min | |
| Denaturation | 98 | 30 sec | 25x |
| Annealing | 59.7 | 30 sec | |
| Elongation | 72 | 30 sec | |
| Final elongation | 72 | 10 min | |
| Storage | 10 | infinite | |

20 µL PCR products were mixed with 4 µL 6x DNA loading dye and 9 µL was separated on a 2% agarose gel [in 1x TAE buffer (40 mM Tris, 20 mM glacial acetic acid, 1 mM EDTA), containing 1x GelRed nucleic acid stain]. 2 µL Quick-Load 100 bp DNA ladder (New England Biolabs, Germany) was used per gel and gels were run at 100 V for 100 min. Gels were documented on an INTAS Sciente Imaging gel documentation system (Supplementary Fig. 7a, summarised in Supplementary Table 7). Apparent *fpvAI*-positive strains were verified by an additional duplex PCR using the primer pairs FpvAI-F/FpvAI-R and PyoS2/ImS2-F and PyoS2/ImS2-R (Supplementary Fig. 7b).

#### Isolation and identification of *P. aeruginosa* strains from human patients

Multidrug-resistant *P. aeruginosa* strains were isolated from anonymised clinical specimens obtained from human patients at the University Hospital Düsseldorf between 2002 and 2020. Isolation and identification were performed using standard diagnostic microbiology procedures, including culturing on selective and non-selective agar media (e.g., cetrimide agar, blood agar). Species were confirmed by using matrix-assisted laser desorption/ionisation-time of flight (MALDI-TOF) mass spectrometry (Vitek MS, bioMérieux, Germany) and antimicrobial susceptibility testing with the semi-automatic VITEK-2 system (bioMérieux, Nürtingen, Germany) and agar-diffusion using EUCAST guidelines (v 12.0, 2020) and breakpoints. For long-term storage, bacterial strains were preserved at −80 °C in Microbank vials (Pro-Lab Diagnostics Inc., Toronto, Canada). For a detailed list of clinical isolates used in this work, see Supplementary Table 7. For the results of antimicrobial susceptibility testing (only available for the core strain collection of *fpvAI*-positive isolates), see Supplementary Table 8.

#### Characterisation of ESBL and carbapenemase genes

All *P. aeruginosa* isolates showing resistance to the carbapenems imipenem/meropenem during antimicrobial susceptibility testing were subjected to multiplex real-time PCR analysis to identify the encoded ESBL or carbapenemase resistance genes (cf. Supplementary Table 7). Strains were plated on MacConkey agar and a single colony was suspended in 20 mM Tris-HCl, 1 mM EDTA, pH 8 (200 µL). Cells were lysed by heating at 95 °C for 10 min, followed by centrifugation (10,000 x g, 10 min, 4 °C). The DNA-containing supernatant (150 µL) was transferred into a new tube and stored at −20 °C for further analyses. The isolated DNA of all *P. aeruginosa* isolates was then analysed in an in-house TaqMan-based multiplex real-time PCR protocol, established for the detection of the most common carbapenemase genes ($bla_{IMP}$, $bla_{VIM-1}$, $bla_{VIM-2}$, $bla_{GIM-1}$, $bla_{GES}$, $bla_{OXA-48}$, $bla_{NDM-1}$ and $bla_{KPC}$)[27-28]. DNA oligonucleotides and fluorescently labelled TaqMan probes used for the detection of β-lactamase resistance genes are summarised in Supplementary Table 11.

#### Isolation of genomic DNA of *P. aeruginosa*

The genomic DNA of *P. aeruginosa* W15Dec1 and 15 MDR clinical isolates for WGS was isolated using the Qiagen Blood & Cell Culture DNA Mini Kit and the protocol for the preparation of Gram-negative bacterial samples. Briefly, the strains were grown overnight in LB medium at 37 °C and an $OD_{600}$ of 2.0 was harvested by centrifugation (6,000 x g, 10 min, 21 °C). The supernatant was discarded and the pellet was thoroughly resuspended in 1 mL buffer B1 (supplemented with 2 µL 100 mg/mL RNase A) by vortexing. 20 µL of lysozyme stock solution (100 mg/mL in water) and 45 µL of Qiagen protease (reconstituted in 1.4 mL water) were added, and the bacterial suspension was incubated for 45 min at 37 °C for lysis. 350 µL of buffer B2 was added and the tube was vortexed for 5 s, followed by an incubation at 50 °C for 30 min. The sample was vortexed again for 10 s and applied to the Qiagen Genomic-Tip 20/G (equilibrated with 1 mL buffer QBT). After complete immersion of the sample into the resin, the 20/G column was washed with buffer QC (4 x 1 mL). Subsequently, the genomic DNA was eluted with prewarmed buffer QF at 50 °C (2 x 1 mL) and collected in a 15 mL tube. DNA was then precipitated by adding 1.4 mL *i*PrOH (0.7 volumes) at 20 °C and carefully inverting the tube 20 times. Immediate pelleting of the DNA by centrifugation (15,000 x g, 20 min, 4 °C) and careful removal of the supernatant yielded a white pellet, which was then washed with ice-cold 70% EtOH (1 mL, kept at –20 °C) by brief vortexing for 2 s and centrifugation (15,000 x g, 30 min, 4 °C). The supernatant was carefully discarded, and the pellet was air-dried for 10 min. Finally, the genomic DNA was resuspended by the addition of 2 mL TE buffer (10 mM Tris-HCl, 1 mM EDTA, pH 8.0) and incubation overnight at 55 °C with shaking at 400 RPM. A final centrifugation step (15,000 x g, 10 min, 4 °C) was applied to separate undissolved particles from reconstituted genomic DNA. The nucleic acid isolates were concentrated in a vacuum centrifuge at 60 °C to the desired concentration, then frozen in liquid nitrogen and finally stored at –20 °C until needed.

#### Whole-genome sequencing and genome assembly

Sequencing library preparation and whole genome sequencing of *P. aeruginosa* genomic DNA was performed by AZENTA/Genewiz (Leipzig, Germany) on an Illumina platform with the NovaSeq 2x150 bp paired-end configuration. A total amount of 2000 ng of genomic DNA of *P. aeruginosa* W15Dec1 and the 15 MDR clinical isolates from the core strain collection (see Fig. 3c) was diluted in TE buffer (10 mM Tris-HCl, 1 mM EDTA, pH 8.0) to a final concentration of 200 ng/µL and a volume of 100 µL was submitted to AZENTA for NGS sequencing. The obtained paired-end raw sequencing data in fastq format were uploaded to the European Galaxy Project webserver (usegalaxy.eu)[29].

Raw paired-end 2×150 bp Illumina reads were first assessed with FastQC to evaluate base quality, adapter content, and sequence composition, and the reports were summarised with MultiQC. Adapter removal, quality trimming, and read filtering were then performed with fastp[30], with parameters adjusted as follows: adapter sequence auto-detection enabled; qualified quality Phred score: 20; unqualified percent limit: 40; minimum read length: 50 bp; polyG tail trimming enabled (Illumina NovaSeq mode); and 3′ end trimming based on quality (window size: 4 bp; mean quality threshold: 20). The processed reads were re-analysed with FastQC, and the results were aggregated using MultiQC to confirm improvements in read quality.

*De novo* assembly was performed with Unicycler[31], which internally employs SPAdes for short-read assembly, Bowtie2 for read alignment, BLAST+ for repeat resolution, and two iterations of polishing with Pilon to remove substitution and indel errors. Normal bridging mode was used and contigs shorter than 300 bp were excluded to reduce assembly artefacts. Assembly quality and completeness were evaluated using QUAST (for contiguity metrics), BUSCO (for single-copy orthologue recovery), and NCBI FCS-GX (for contamination screening). Read alignments were processed with SAMtools, and coverage across the assembled genome was visualised in JBrowse2. Annotation was conducted with Bakta[32] using a minimum contig size threshold of 300 bp, the Bakta database v5.1, and the AMRFinderPlus database v3.12, with the genus specified as *Pseudomonas* to optimise functional annotation. Genome assemblies and annotations were visualised and customised using Artemis[33] (Wellcome Sanger Institute).

Final genome assemblies (FASTA format) were deposited in the NCBI genome repository under the BioProject PRJNA1336694 and annotated using NCBI's PGAP pipeline[34]. The following DDBJ/ENA/GenBank accession numbers were assigned: W15Dec1 (JBRLAX010000000), M2 (JBRLBM010000000), M7 (JBRLBL010000000), M10 (JBRLBK010000000), M71 (JBRLBJ010000000), M74 (JBRLBI010000000), M132 (JBRLBH010000000), M134 (JBRLBG010000000), M189 (JBRLBF010000000), M190 (JBRLBE010000000), M191 (JBRLBD010000000), M210 (JBRLBC010000000), M294 (JBRLBB010000000), M303 (JBRLBA010000000), M315 (JBRLAZ010000000), and M324 (JBRLAY010000000).

#### Genome mining for serotypes, defects in LPS biogenesis, and resistance genes

Assembled genomes of *P. aeruginosa* isolates from the core strain collection of *fpvAI*-positive strains from the University Hospital in Düsseldorf (cf. Fig. 3c) were screened for certain genes of interest and single nucleotide polymorphisms by using the local NCBI blast+ algorithms (v2.17.0) tBLASTn[35] and BLASTp[36]. Therefore, the assembled genomes were searched against reference sequences of all proteins of interest (LPS biosynthesis genes, pyocins, and antibiotic resistance genes, see supplementary MOESM1_ESM word file), retrieved from reference genomes of *P. aeruginosa* strains, and the loci were subsequently mapped against the PGAP annotation files using Python (v3.13.7) to identify overlaps between blast search results and annotated features (protein identifiers and locus_tags, see supplementary MOESM2_ESM excel file):

- **strain PAO1** (NC_002516.2, IATS serogroup O5; source for all reference sequences except those specified in the following paragraph)[19],
- **strain IATS:O6** (GenBank AF035937.1, reference for IATS serogroup O6)[37],
- **strain PA103** (GenBank AF147795.1, reference for IATS serogroup O11)[38],
- **strain UCBPP-PA14** (NC_008463.1, source for exoU, accession number WP_003134060.1)
- **strain DSM 50071** (GenBank CP012001.1, source for pyocin S1, accession number AKO87955.1)
- **strain P12** (GenBank X77996.1, source for pyocin S3, accession number CAA54958.1)
- **strain 105738** (LOHK01000018.1, source for pyocin S6, accession number KYO98147.1)
- **strain BWHPSA018** (GenBank AXQK01000000.1, source for pyocin S7, accession number ERW54179.1)
- **strain BL04** (GenBank AXPW00000000.1, source for pyocin S9, accession number ERV82772.1)
- **strain PABL056** (Genome Assembly GCF_000290555.2, source for pyocin S10, accession number WP_025991911.1)
- **strain BWHPSA019** (AXQJ01000003.1, source for pyocin SD1, accession number WP_023116694.1)
- **strain MSH-10** (NZ_KE138672.1, source for pyocin SD2, gene not annotated; accession number of the cognate immunity protein WP_003101816.1)
- **strain PA7** (NC_009656.1, source for pyocin SD3, accession number ABR82489.1)
- **strain WH-SGI-V-07287** (LLUM01000014.1, source for pyocin SX1, accession number KSR45004.2)
- **strain 2_1_26** (GenBank ACWU01000201.1, source for pyocin SX2, accession number EHF11209.1)
- **strain BWHPSA046** (GenBank AZZH01000005.1, source for pyocin G, accession number ETV05907.1)
- **strain PAF41-2** (GenBank D12705.1, source for pyocin AP41, accession number BAA02196.1)
- **strain 7NSK2** (source for FpvAIIa, accession number AF537095.2)
- **strain ATCC 27853** [NZ_CP015117.1, source for FpvAIIb, accession number WP_016852868.1 (wrong annotation)]
- **strain 59.20** (NZ_NSUN00000000.1, source for FpvAIII, accession number AAN62912.1)

Besides the CPA and OSA biosynthesis genes listed in Supplementary Fig. 8, the following LPS biosynthesis genes were screened for anomalies that would explain ImS2-independent resistance to PyoS2 Y640C proteins:
Lipid A and LPS biosynthesis: *migA*, *dnpA*, *waaA*, *waaC*, *waaF*, *waaL*, *waaP*, *waaG*, *wapG*, *wapH*, *wapO*, *wapP*, *wapR*, *galU*, *algC*, *gmhA*, *gmhB*, *rfaE/hldE*, *rfaD/hldD*, *rmlA–D*, *ytfN*, *pagL*, *arnT*, *lpxA–D*, *lpxH*, *lpxK*.
LPS assembly: *wzz2*.
Complete IATS O6 cluster: *wbpO*, *wbpP*, *wbpQ*, *wzx*, *wbpR*, *wbpS*, *wbpT*, *wbpU*, *wbpV*, *wbpL*, *wbpM*.
Complete IATS O11 cluster: *wzz*, *wzx*, *wzy*, *wbjA*, *wbjB*, *wbjC*, *wbjD*, *wbjE*, *wbjF*, *wbpL*, *wbpM*.
LPS transport: *lptA–G*.
However, no common anomalies were detected in the loci mentioned above.

Multiple sequence alignments of nucleotide and protein sequences were generated with Clustal Omega[39] and were visualised and edited with Jalview[40]. Prediction of O serogroups based on bacterial genomes was performed with the web version of the *Pseudomonas aeruginosa* serotyper 1.0 (PAst 1.0)[41-42]. Reference sequences of β-lactamase resistance genes were retrieved from the Beta-Lactamase Database (BLDB)[43].

#### Plasmid nicking assay (*in vitro* DNase activity)

The plasmid nicking assay[44] was used to validate the DNase activity of PyoS2 mutants and PyoS2 conjugates relative to wild-type PyoS2 in the presence or absence of the inhibitory immunity protein ImS2. For the characterisation of PyoS2 DNase activity without ImS2 inhibition, PyoS2 derivatives were thawed on ice and diluted to 40 µM in plasmid nicking buffer (25 mM Tris-HCl, 150 mM NaCl, 10 mM $MgCl_2$, pH 8.0). For DNase activity in the presence of the innate PyoS2 inhibitor ImS2, both proteins were diluted in the same tube to 40 µM and 42 µM, respectively, and incubated at RT for 20 min to allow complex formation and incorporation of the $Mg^{2+}$ cofactor. pET21a(+) plasmid DNA (177.3 ng/µL, stock solution in water) was used as a DNA substrate for the nuclease bacteriocin PyoS2.

In a 1.5 mL reaction tube, pET21a(+) plasmid DNA (assay concentration 26.7 ng/µL), plasmid nicking buffer and PyoS2 (assay concentration 5 µM) or PyoS2/ImS2 (assay concentration 5 µM/5.25 µM) were mixed to give a final volume of 90 µL. The plasmid nicking was carried out at 30 °C (400 RPM). For the kinetic experiment, 7.5 µL of the reaction mixture was taken after 2, 5, 10, 20, 30, 40, 50, 60, 70, 80, and 90 min and immediately quenched by adding the reaction mixture to a solution of 0.25 µL 1 M EDTA and 1.25 µL 6x DNA loading dye. A 0 min sample was prepared separately by mixing DNA, plasmid nicking buffer, and quenching buffer in advance and finally adding the enzyme to a final concentration of 5 µM PyoS2 (optional: 5.25 µM ImS2). pET21a(+) DNA without any enzyme served as a negative control. Alternatively, the same setup was used for long-term monitoring of DNA degradation. For this long-term monitoring, only a single sample was taken after 15 h.

For the analysis of DNase-catalysed pET21a(+) DNA digest, 2 µL of 1 kb DNA ladder and 9 µL quenched reaction mixture (200 ng/DNA per well) for each time point were loaded onto a 0.8% agarose gel (with 1x GelRed stain) in 1x TAE buffer and the gel was run for 40 min at 180 V. Gels were subsequently analysed and documented on an INTAS Science Imaging gel documentation system for further densitometric analyses.

#### Densitometric analysis of DNase activity

Photographs of DNase agarose gels were analysed with Fiji ImageJ2[45] to quantify the relative amount of nicked, linear, and supercoiled DNA from the measured pixel density (densitometry). The average pixel density of unloaded wells was subtracted as background, and the corrected gels were further analysed using the gel analyser tool. A uniform "rectangle" was drawn around each form of DNA individually and the lanes were plotted against pixels. Using the "straight" line tool, a baseline was introduced manually for each peak and the integral was calculated as pixel density with the "wand (tracing) tool". The obtained pixel densities Y for each DNA form and time point were then normalised to the sum of nicked (N), linear (L), and supercoiled (S) DNA at the $t$=0 time point ($t$=0 equals 100%, equation 1) and their relative percentage for each time point $t_x$ plotted against the time.

$$Y_{x,norm}\,[\%] = 100 \cdot \frac{Y_x}{Y_{N,0} + Y_{L,0} + Y_{S,0}} \qquad \text{(eq. 1)}$$

| | | |
|---|---|---|
| $Y_x$ | = | Pixel density at time point x |
| $Y_{N,0}$ | = | Pixel density of nicked DNA at $t$=0 |
| $Y_{L,0}$ | = | Pixel density of linear DNA at $t$=0 |
| $Y_{S,0}$ | = | Pixel density of supercoiled DNA at $t$=0 |
| $Y_{x,norm}$ | = | Relative percentage of nicked, linear, or supercoiled DNA at $t_x$ (relative to $t$=0) |

To quantitatively describe the degradation of supercoiled DNA over time, each dataset was independently fitted to a stretched exponential decay model (for highly active DNase mutants, equation 2.1) or to a linear model (for DNase mutants with residual activity, equation 3.1). Therefore, the relative portions of supercoiled DNA were extracted for every time point via densitometric analysis from the corresponding agarose gel of the time-course experiment. Subsequently, the plotted data (relative $DNA_{supercoiled}$ portion vs. time) were analysed according to equation 2.1 or equation 3.1. Furthermore, Pearson correlation analyses (equation 4) and permutation analyses (equation 5) were performed.

#### Stretched exponential decay (Kohlrausch–Williams–Watts function)

Each dataset (see below) was independently fitted to a stretched exponential decay model of the form

$$y(t) = y_0 \cdot e^{-(k \cdot t)^b} \qquad \text{(eq. 2.1)}$$

with

$$t_{\frac{1}{2}} = \frac{[\ln(2)]^{1/b}}{k} \qquad \text{(eq. 2.2)}$$

| | | |
|---|---|---|
| $b$ | = | stretching exponent |
| $y$ | = | measured variable [supercoiled DNA portion at time x] |
| $y_0$ | = | initial amplitude at $t$=0 [% of supercoiled DNA at $t$=0] |
| $k$ | = | decay rate |
| $t_{1/2}$ | = | half-life of supercoiled DNA |

using nonlinear least-squares regression implemented in R (nlsLM, minpack.lm package)[46]. The initial value $y_0$, decay rate parameter k, and stretching exponent b were estimated for each dataset, and 95% confidence intervals were derived when the fit permitted. Goodness-of-fit was assessed via the coefficient of determination ($R^2$), residual standard error (RSE), and visual inspection of residuals. The half-life $t_{1/2}$, the time at which the fitted signal decayed to 50% of $y_0$, was estimated numerically by linear interpolation according to equation 2.2. For each fit, three diagnostic plots were generated: (i) the raw data with fitted curve, (ii) residuals versus time, and (iii) a log-linear transformation of the observed signal to assess conformity to exponential behaviour.

The exponential decay analysis was performed for the following datasets of PyoS2 mutants and conjugates with an apparent exponential decrease of supercoiled DNA species (high DNase activity):

PyoS2 (toxin only): wild-type; Y640C; Q570C-AF488; K602C-AF488; Y640C-AF350; Y640C-AF488; Y640C-AF594; C-Cys-AF488.

PyoS2 (+ImS2): Y640C-AF350 + ImS2; Y640C-AF488 + ImS2; Y640C-AF594 + ImS2.

Values of supercoiled DNA half-life and exponential decay fit parameters are summarised in Supplementary Table 4. Graphical analyses are shown in Supplementary Fig. 4.

#### Linear regression

Each dataset (see below) was independently analysed using linear regression according to the following model:

$$y(t) = b \cdot t + y_0 \qquad \text{(eq. 3.1)}$$

with

$$t_{\frac{1}{2}} = \frac{\frac{y_0}{2} - y(0)}{b} \qquad \text{(eq. 3.2)}$$

| | | |
|---|---|---|
| $b$ | = | slope; rate of change |
| $y$ | = | measured variable (supercoiled DNA portion at time x) |
| $y_0$ | = | y-intercept (fitted parameter) |
| $t_{1/2}$ | = | half-life of supercoiled DNA |
| $y(0)$ | = | supercoiled DNA portion at $t$=0 min (experimental). |

with time as the independent variable and the relative portion of supercoiled DNA as the dependent variable. The analysis was performed in R[46] using the *lm()* function, and summary statistics were extracted using the *broom* package. The initial signal intensity $y_0$ was defined as the value of the first time point in each dataset. The half-life $t_{1/2}$, defined as the time at which the fitted linear model predicted a signal of $y_0/2$, was calculated algebraically from the model intercept and slope according to equation 3.2. Goodness-of-fit was assessed using the coefficient of determination ($R^2$), adjusted $R^2$, standard error, p-value for the slope, and the F-statistic. Diagnostic plots were generated for each dataset, including (i) the fitted regression line overlaid on the experimental data, and (ii) the residuals plotted against the fitted values.

The linear regression analysis was performed for the following datasets of PyoS2 mutants and conjugates with an apparent linear decrease of supercoiled DNA species:

PyoS2 (toxin only): H685A.

PyoS2 (+ImS2): Y640C + ImS2.

Values of supercoiled DNA half-life and fit parameters are summarised in Supplementary Table 6. Graphical analyses are shown in Supplementary Fig. 5.

#### Pearson correlation analysis

To assess whether the observed concurrent decline of supercoiled, linear, and nicked plasmid DNA species over time was attributable to assay-related degradation (systematic error) rather than DNase-mediated cleavage, a Pearson correlation analysis was conducted using OriginLab.

In theory, enzymatic digestion of supercoiled plasmid DNA by an active PyoS2 DNase is expected to initially yield an increase in linear DNA – the intermediate between supercoiled and fully degraded products – before its eventual degradation into smaller fragments. In contrast, DNase-inactive protein variants showing a synchronous decline across all DNA forms (without a transient increase in linear DNA) may reflect assay-related artefacts such as sample degradation or binding of DNA to the plastic of the reaction tubes.

Pearson's correlation coefficient (r) was used to measure the strength and direction of the linear relationship between the abundance profiles of the different DNA species over time. The Pearson correlation coefficient is defined as:

$$r = \frac{\sum_{i=1}^{n}(x_i - \bar{x})(y_i - \bar{y})}{\sqrt{\sum_{i=1}^{n}(x_i - \bar{x})^2}\sqrt{\sum_{i=1}^{n}(y_i - \bar{y})^2}} \quad \text{(eq. 4)}$$

| | | |
|---|---|---|
| $x_i$ and $y_i$ | = | values of two variables (e.g. intensity of supercoiled and nicked DNA) at time point $i$ |
| $\bar{x}$ and $\bar{y}$ | = | mean values of respective datasets |
| $n$ | = | number of time points measured. |

In this context, a strong positive correlation (r close to +1) between all three DNA species suggests a shared mode of signal loss, such as systematic assay degradation rather than enzymatic action. Conversely, weak or negative correlations may indicate true enzymatic cleavage behaviour, where one species (e.g. supercoiled) declines while another (e.g. linear) transiently increases.

This analysis helped differentiate between protein variants with genuine DNase activity and those whose apparent effects on DNA integrity arose from assay-related phenomena. Pearson coefficients and correlation graphs are summarised in Supplementary Fig. 3.

#### Permutation analysis

To assess whether the slow increase in linear plasmid DNA over time – observed in time-resolved plasmid nicking assays with low-activity DNase mutants – was statistically significant, a permutation analysis was performed using R with 10,000 random permutations and a 95% confidence interval. Permutation testing is a non-parametric method that evaluates significance by comparing the observed test statistic (here, the coefficient of determination $R^2$ from a linear regression of linear DNA vs. time) against a distribution of $R^2$ values generated from randomly shuffled data under the null hypothesis. This approach avoids assumptions about data normality and is well-suited for small or irregular biological datasets.

The test statistic used was the coefficient of determination ($R^2$), calculated as:

$$R^2 = 1 - \frac{\sum_{i=1}^{n}(y_i - \hat{y})^2}{\sum_{i=1}^{n}(y_i - \bar{y})^2} \quad \text{(eq. 5)}$$

| | | |
|---|---|---|
| $y_i$ | = | observed linear DNA at time point $i$ |
| $\hat{y}_i$ | = | predicted value from regression |
| $\bar{y}$ | = | mean values of all observed values |
| $n$ | = | number of time points measured. |

The empirical p-value was determined as the proportion of permuted $R^2$ values greater than or equal to the observed $R^2$. Three plots were used to interpret the regression output: i) Histogram of permuted $R^2$ values [distribution of $R^2$ under the null hypothesis], ii) residuals vs. time [assessment of model fit; randomly scattered residuals suggest homoscedasticity and linearity], and iii) observed vs. predicted values [agreement between actual and fitted values]. Together, the $R^2$ value and permutation-derived p-value provided statistical validation for a genuine, time-dependent increase in linear DNA, distinguishing low-level DNase activity from random variation or assay noise. All graphs and fit parameters generated during the permutation analysis can be found in Supplementary Fig. 5 and Supplementary Table 5.

#### Agar spot overlay assay/plate killing assay

The plate killing assay[47] was used to assess the cytotoxic potential of pyocin S2 and its conjugates against selected strains of *P. aeruginosa*. First, LB medium was inoculated with a single colony of the desired *P. aeruginosa* strain and incubated overnight at 37 °C on a rotary shaker. 20 mL of LB medium was then inoculated with 200 µL of the overnight culture and incubated at 37 °C for 2.5–3 h until visible turbidity was obtained ($OD_{600}$ not measured). 5 mL of this exponential day culture was mixed with 45 mL liquid 0.7% LB (kept at 50 °C until needed) and 7 mL of this culture was immediately poured over a 1.4% LB agar plate to cover the 1.4% LB agar base with a layer of immobilised cells. After hardening at room temperature, the plates were briefly dried under the safety cabinet. 5 µL of protein dilutions (10 µM–57 pM) in sterile-filtered dialysis buffer (25 mM Tris-HCl, 150 mM NaCl, pH 8) were spotted onto the surface. After evaporating the excess liquid under the safety cabinet, the plates were incubated overnight at 37 °C, and antimicrobial activity was subsequently documented by photography. Clear spots of inhibited bacterial growth indicate cytotoxic activity related to PyoS2 treatment.

##### Turbidity assay/liquid growth inhibition assay

The liquid growth inhibition assay was used to validate PyoS2 killing and killing activity of its conjugates against *P. aeruginosa* PAO1 and W15Dec1 in liquid media. First, LB medium was inoculated with a single colony of the desired *P. aeruginosa* strain and incubated overnight at 37 °C on a rotary shaker. 20 mL of LB medium was then inoculated with overnight cultures of *P. aeruginosa* PAO1 or W15Dec1 to an $OD_{600}$ of 0.05 and distributed to a 24-well plate (ThermoScientific, Nunclon Delta Surface, Catalogue No. 142475, sterile, flat-well bottom, transparent with lid) with 2 mL per well. The plate was incubated in a Molecular Devices SpectraMax ID3 plate reader at 37 °C with medium shaking, and the optical density was monitored at 600 nm in intervals of 2 min (for instrument settings see below). For strain W15Dec1, 30 µL of filter-sterilised dialysis buffer, 30 µL of 50 µM PyoS2 (750 nM final concentration), or 30 µL of 50 µM PyoS2 Y640C AF350 (750 nM final concentration) were added after 1:49 h. For strain PAO1, the same proteins and volumes were added after 2:51 h. Monitoring of triplicate samples was continued until a total incubation time of 1000 min was reached. Dialysis buffer was used as a negative control and LB medium was subtracted from all samples as background.

<u>The following data record parameters were used:</u>
Temperature: 37 °C; Mode: Kinetic [Time: 15:00 h, Interval: 02:00 min, Reads: 451, Min. interval: 01:57 min, Absorbance: 600 nm]; Plate type: 24 well standard clear bottom (landscape, lidded, 23 mm height); Additional settings: shake before first read (5 sec, medium speed, orbital), shake between reads (100 sec, medium speed, orbital), speed read (on), read order (row), show optimiser (on). Before the first read, plate optimisation was performed (ca. 9 min) with the filled plate already in the instrument at 37 °C.

##### Protein sequences of PyoS2 and ImS2

<u>Protein sequence of wild-type PyoS2 from *P. aeruginosa* PAO1:</u>

```
MAVNDYEPGSMVITHVQGGGRDIIQYIPARSSYGTPPFVPPGPSPYVGTGMQEYRKLRSTLDKSHSELKKNLKNETLKEVDELKSEAGLPGKAVSANDIRDEK
SIVDALMDAKAKSLKAIEDRPANLYTASDFPQKSESMYQSQLLASRKFYGEFLDRHMSELAKAYSADIYKAQIAILKQTSQELENKARSLEAEAQRAAAEVEA
DYKARKANVEKKVQSELDQAGNALPQLTNPTPEQWLERATQLVTQAIANKKKLQTANNALIAKAPNALEKQKATYNADLLVDEIASLQARLDKLNAETARRKE
IARQAAIRAANTYAMPANGSVVATAAGRGLIQVAQGAASLAQAISDAIAVLGRVLASAPSVMAVGFASLTYSSRTAEQWQDQTPDSVRYALGMDAAKLGLPPS
VNLNAVAKASGTVDLPMRLTNEARGNTTTLSVVSTDGVSVPKAVPVRMAAYNATTGLYEVTVPSTTAEAPPLILTWTPASPPGNQNPSSTTPVVPKPVPVYEG
ATLTPVKATPETYPGVITLPEDLIIGFPADSGIKPIYVMFRDPRDVPGAATGKGQPVSGNWLGAASQGEGAPIPSQIADKLRGKTFKNWRDFREQFWIAVAND
PELSKQFNPGSLAVMRDGGAPYVRESEQAGGRIKIEIHHKVRIADGGGVYNMGNLVAVTPKRHIEIHKGGK
```

<u>Protein sequence of ImS2-$His_6$ from *P. aeruginosa* PAO1 with artificial $His_6$-tag:</u>

```
MKSKISEYTEKEFLEFVKDIYTNNKKKFPTEESHIQAVLEFKKLTEHPSGSDLLYYPNENREDSPAGVVKEVKEWRASKGLPGFKAGLEHHHHHH
```
